# Neuropeptide CRH prevents premature differentiation of OPCs following CNS injury and in early postnatal development

**DOI:** 10.1101/2025.02.14.638041

**Authors:** Clemens Ries, Tibor Stark, Benoit Boulat, Torben Ruhwedel, Jan Philipp Delling, Antonio Miralles Infante, Julia T. von Poblotzki, Alessandro Ulivi, Iven-Alex von Mücke-Heim, Simon Chang, Kenji Sakimura, Keiichi Itoi, Dennis B. Nestvogel, Alessio Attardo, Michael Czisch, Klaus-Armin Nave, Wiebke Möbius, Leda Dimou, Jan M. Deussing

## Abstract

The role of neuropeptides and their receptors in oligodendrocyte progenitor cells (OPCs) has largely been overlooked so far. Here, we describe a new subpopulation of corticotropin-releasing hormone (CRH)-expressing OPCs that aggregate around acute brain injuries and exhibit an elevated capacity to differentiate into myelinating oligodendrocytes (OLs). We found that CRH expression in OPCs is rapidly induced *de novo* as a transient response within the first 72 hours after injury. As target cells, we identified CRH receptor type 1 (CRHR1)-expressing OPCs which show a decreased differentiation velocity. We demonstrate that CRH/CRHR1 system inactivation increases the speed of OL generation compromising the long-term survival of OLs after acute injury. Furthermore, we prove that a CRH/CRHR1 system deficiency under non-injury conditions leads to increased early postnatal oligodendrogenesis and alterations in adult myelination. Altogether, we show that OPC-derived CRH not only actively influences the injury environment through the interaction with CRHR1-expressing OPCs, but also identify the G-protein coupled receptor CRHR1 as a critical modulator of oligodendrogenesis at early postnatal stages with lasting effects on adult myelination.

## Introduction

Oligodendrocyte progenitor cells (OPCs), are the main source of myelinating oligodendrocytes (OLs) and have been widely accepted as the fourth major glial cell type of the central nervous system (CNS) (Jakel and Dimou, 2017; Nishiyama et al., 2021; Raff et al., 1983). Upon acute CNS injury, OPCs proliferate, migrate towards the injury site and differentiate into OLs (Dimou et al., 2008; Hughes et al., 2013; Simon et al., 2011; Vigano et al., 2016; von Streitberg et al., 2021). In regeneration, but also in development and adulthood, oligodendrogenesis is an inefficient process characterized by an overproduction of premyelinating OLs, of which only a fraction reaches the mature stage (Barres et al., 1992; Hughes et al., 2018; Trapp et al., 1997). While the exact mechanisms of differentiation initiation and myelination are still not fully understood, they have already been shown to be influenced by many different factors including neuronal activity and G-protein coupled receptor (GPCR) signaling, e.g., by the kappa opioid receptor and its neuropeptide ligand dynorphin (Gibson et al., 2014; Hughes *et al*., 2018; Hughes and Stockton, 2021; Osso et al., 2021).

Neuropeptides, such as dynorphin, constitute a diverse and heterogeneous group of signaling molecules that target a wide range of structures and biological functions. Stored in large dense core vesicles, they signal via a mode known as volume transmission which distinguishes these neuromodulators from classical neurotransmitters. Accordingly, locally released neuropeptides can have biological effects in a micrometer range (Ozcete et al., 2024; van den Pol, 2012). Neuropeptides are typically released in large quantities and exert their biological functions through binding and signaling via GPCRs present on target cells. One of the best-studied neuropeptides is the corticotropin-releasing hormone (CRH) which is expressed in neurons throughout the brain. Together with its high affinity CRH receptor type 1 (CRHR1), CRH orchestrates the neuroendocrine, autonomic and behavioral stress response (Deussing and Chen, 2018; Joels and Baram, 2009). While the CRH/CRHR1 system has anecdotally been reported to affect the brain’s reaction to physical damage, its role in injury-connected regenerative processes is unknown (de la Tremblaye et al., 2017; Stevens et al., 2003).

In this study, we identify a CRH/CRHR1 system in OPCs that is activated in response to injury. We show that a subpopulation of OCPs rapidly induces CRH expression following acute injury, which acts through CRHR1 on a separate OPC population to modulate OL generation. Furthermore, we demonstrate that the CRH/CRHR1 system influences the dynamics of oligodendrogenesis not only following injury but also under non-injury conditions during early postnatal and adult stages.

## Results

### *De novo* expression of CRH in a subpopulation of OPCs in response to acute brain injury

To study the distribution and connectivity of CRH-expressing neurons in the murine brain, stereotaxic injections of fluorescent retro-beads were performed in the ventral tegmental area (VTA) of *CRH-Cre::Ai9* reporter mice, in which CRH expression is reported via tdTomato (**Fig. 1a, b, Table S1**). Analyzing injected brains, we consistently observed an aggregation of tdTomato^+^ cells with non-neuronal morphology in close proximity to the injection site (**Fig. 1c**). Since the involvement of CRH in the reaction to brain damage was largely unknown, this intriguing observation demanded further investigation. First, we assessed whether *de novo* CRH expression upon acute injury represents a global phenomenon by inflicting injuries in *CRH-Cre::Ai9* mice in: i) prefrontal cortex, ii) striatum, and iii) midbrain (MB). In all regions, the appearance of tdTomato^+^ cells was observed 3 days post injury (dpi) (**Fig. 1d**). To specify the identity of the newly appearing tdTomato^+^ cells, immuno stainings for different glial (GFAP, Iba1 and NG2) and neuronal (NeuN) markers were performed (**Fig. 1e, Fig. S1a-c**). Non-neuronal tdTomato^+^ cells around the injury site only showed co-localization with the OPC marker NG2, specifying these cells as OPCs (**Fig. 1e**). Using the intersectional reporter mouse line *CRH-FlpO::NG2-CreERT2::Ai65* in which co-expression of CRH and NG2 triggers the expression of tdTomato in a FlpO- and Cre-dependent manner, CRH expression in OPCs was confirmed (**Fig. 1f,g, Table S1**). To have a direct measure of CRH expression upon acute injury, we used combined immuno staining against CRH and the OPC-specific marker PDGFRα showing clear co-localization in several cells surrounding the injury site (**Fig. 1h,h’**). Characteristic for a neuropeptide, CRH was localized in vesicular structures throughout the PDGFRα^+^ cells. In summary, we demonstrate that OPCs express the neuropeptide CRH as a reaction to acute injury.

**Fig. 1:**
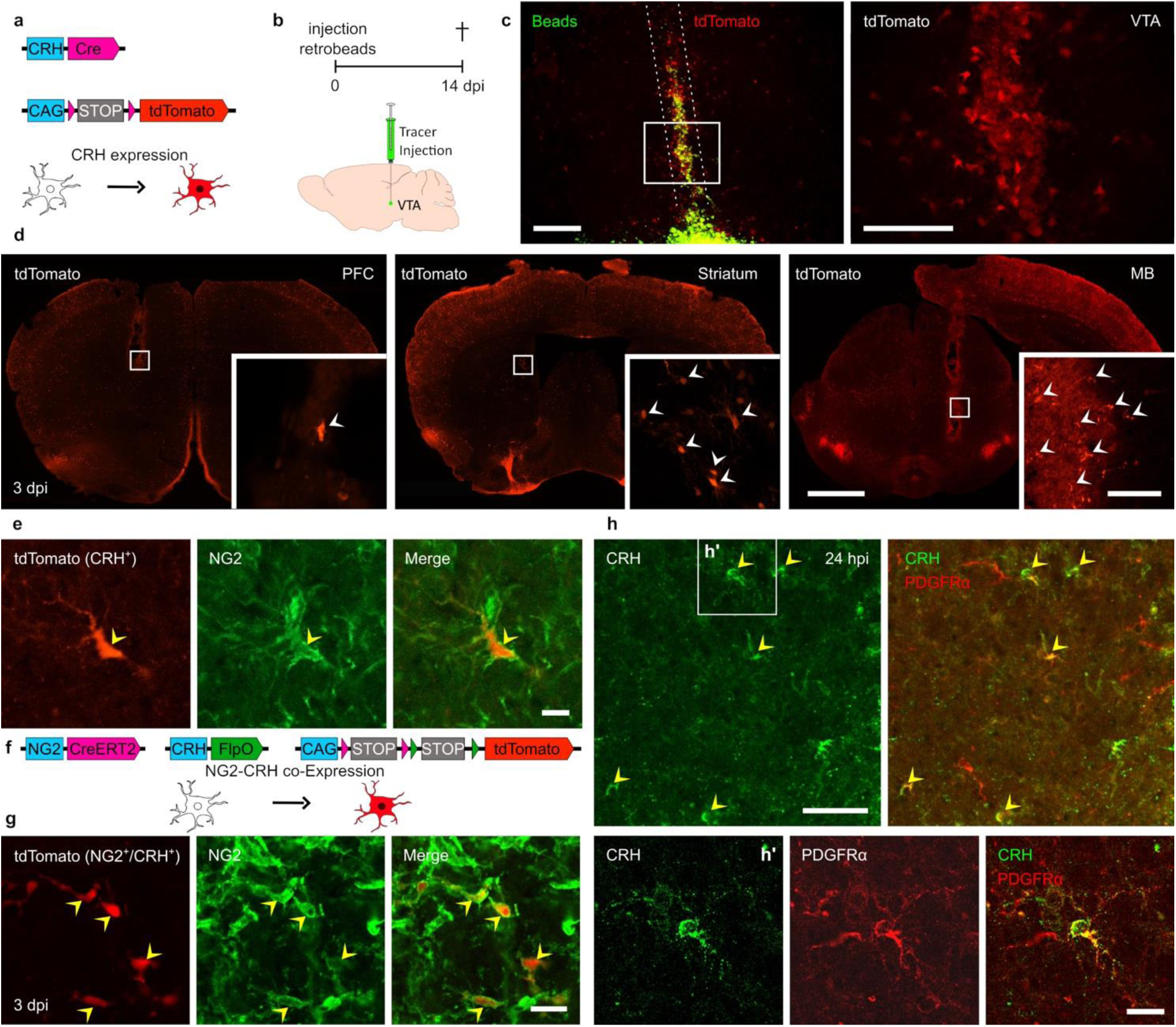
Identification of CRH-expressing OPC subpopulation. **a**, *CRH-Cre::Ai9* reporter mouse model. **b**, Experimental setup and injection site. **c**, Injection site of fluorescent beads and needle tract (framed by white dotted lines) with aggregation of tdTomato^+^ cells around the injury site. Scale bars, 100 µm. **d**, Aggregation of CRH-expressing (tdTomato^+^) cells around injury sites in PFC, striatum and MB in *CRH-Cre::Ai9* mice. Scale bars, 1000 µm (overview), 50 µm (close up). **e**, Immuno staining for NG2 at the injury site in *CRH-Cre::Ai9* mice. Scale bar, 10 µm. **f**, Intersectional reporter mouse line *CRH-FlpO::NG2-CreERT2::Ai65* for monitoring CRH/NG2 co-expression. **g**, Confocal images of NG2/CRH co-expressing (tdTomato^+^) cells at injury site at 3 dpi stained for NG2. Scale bar, 20 µm. **h**, Confocal image of combined CRH/PDGFRα staining at 24 hpi. Scale bar, 50 µm. **h’**, Magnification of CRH/PDGFRα co-expressing cell. Scale bar, 20 µm. For all images: White arrowheads indicate cells or structures. Yellow arrowheads indicate co-localization of two markers.

### CRH-expressing OLCs exhibit a strong proliferative response

To interrogate the population dynamics of CRH-expressing oligodendrocyte lineage cells (OLCs), we first focused on their proliferative capacity and analyzed changes in their cell number after injury. To analyze short- and long-term changes, *CRH-Cre::Ai9* mice were subjected to stab wound injury in the midbrain and sacrificed at 1, 2, 3, 7, 14, 23, 69 or 128 dpi (**Fig. 2a**). tdTomato^+^/Olig2^+^ cells were quantified in a 300 µm radius around the wound center with 50-100 µm medio-lateral resolution (**Fig. 2b,c, Methods S1a-c**). Quantification in the whole area revealed a considerable increase of tdTomato^+^/Olig2^+^ cells between 2 (14 ± 8.13/ mm^2^) and 7 dpi (168 ± 27.85/ mm^2^) followed by a significant decrease until 128 dpi (55 ± 6.56 cells/ mm^2^) (**Fig. 2c**). Also, within the first 50 µm around the wound center, dynamics were comparable with a considerably higher density (614 ± 87.62 cells/mm^2^) **(Fig. S2a).** The medio-lateral distribution of CRH-expressing OLCs over time, depicted in the heatmap, shows their initial appearance within the entire area of analysis (0-300 µm distance) with a progressive inward movement over time (**Fig. 2c**). Because of the well described proliferative reaction of OPCs following insult (von Streitberg *et al*., 2021), we investigated whether cell division following CRH expression contributed to the increase in total cell number. Indeed, tdTomato^+^/Olig2^+^ cells did not only show *de novo* expression of CRH but also co-expression of the proliferation marker Ki67 (**Fig. 2d**). Quantification revealed that, while at 2 dpi the vast majority of Olig2^+^/tdTomato^+^ cells were Ki67^+^ (98 ± 1.75%), the proportion of co-expressing cells dropped significantly until 7 dpi (18 ± 2.31%) (**Fig. 2e, Fig. S2b**).

**Fig. 2:**
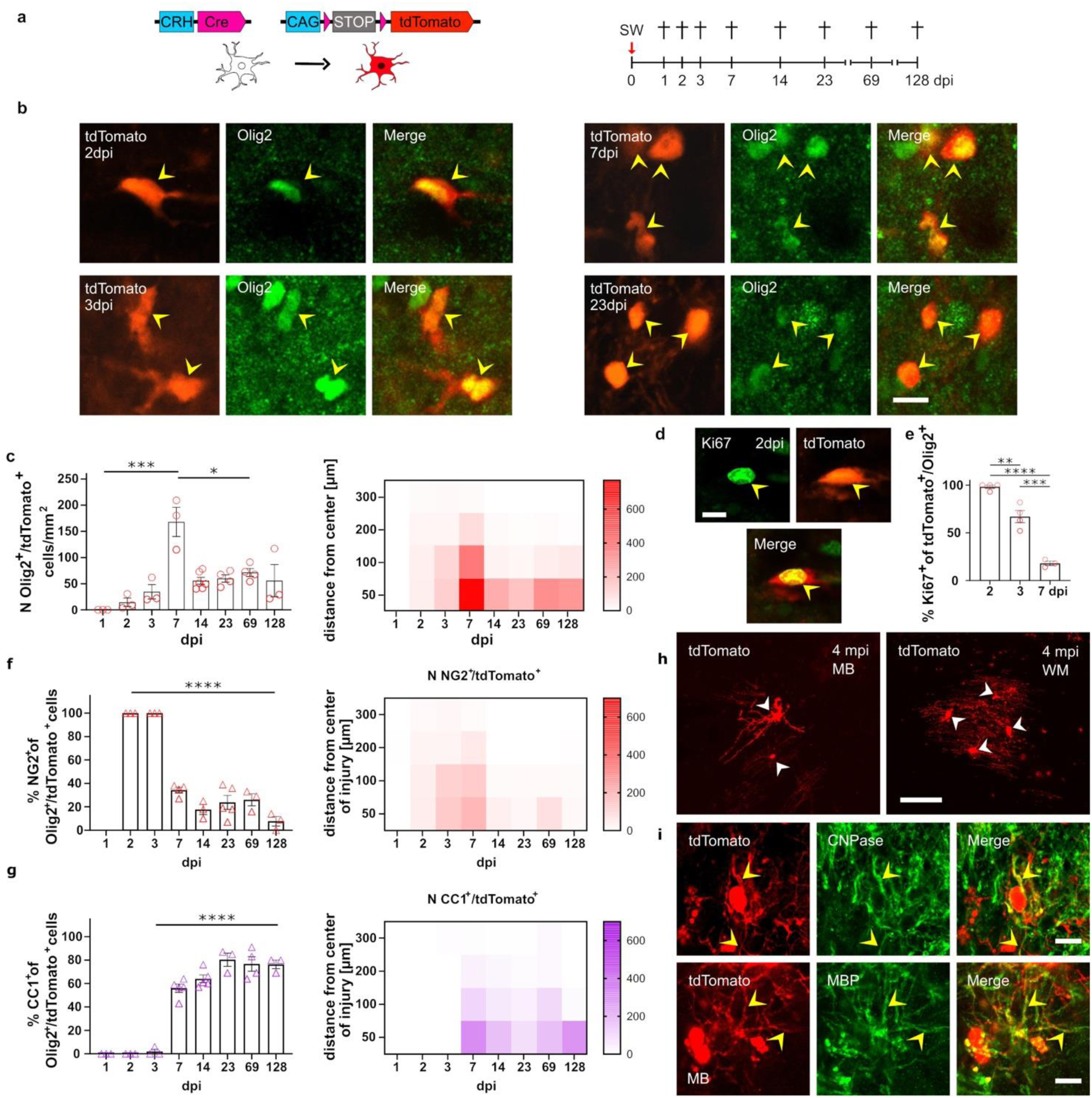
CRH^+^ OLCs proliferate and generate myelinating OLs following injury. **a**, *CRH-Cre::Ai9* mouse model and experimental setup. **b**, Representative confocal images of *CRH-Cre::Ai9* mice at 2, 3, 7 and 23 dpi stained for Olig2. Scale bar, 10 µm. **c**, Quantification of Olig2^+^/tdTomato^+^ cells at ± 300 µm around the injury site (One-way ANOVA: F_(7, 21)_= 11.37, p < 0.0001). 1-7 dpi, (p < 0.001, 95% C.I. = -239,7, -96,35. 7-128 dpi, p = 0.007, 95% C.I. = 40.55, 183.9) and heat map showing spatio-temporal appearance of Olig2^+^/tdTomato^+^ cells. **d**, Confocal image of Ki67 staining at 2 dpi. Scale bar, 10 µm. **e**, Quantification of Ki67^+^/tdTomato^+^/Olig2^+^ of all tdTomato^+^/Olig2^+^. 2-7 dpi, One-way-ANOVA: F_(2, 8)_ = 81.16, p < 0.0001. **f**, Left, percentage of NG2^+^/Olig2^+^/tdTomato^+^ cells out of all Olig2^+^/tdTomato^+^ cells. 3-7 dpi, One-way ANOVA: F_(6, 17)_= 68.5, p < 0.0001. Right, Number of NG2^+^/tdTomato^+^ cells in spatio-temporal resolution. n_TP_ = 3-6 mice. **g**, Left, percentage of CC1^+^/Olig2^+^/tdTomato^+^ cells of all Olig2^+^/tdTomato^+^ cells. 3-7 dpi, One-way ANOVA: F_(6, 17)_= 68.5, p < 0.0001. Right, Number of CC1^+^/tdTomato^+^ cells in spatio-temporal resolution. n_TP_ = 3-6 mice. **h**, Morphologies of tdTomato^+^ cells at 4 mpi in MB (left) and WM (right). Scale bar, 50 µm. **i**, Confocal images of anti-CNPase and anti-MBP staining showing co-localization with tdTomato^+^ cells. Scale bar, 10 µm. White arrowheads: cells or structures. Yellow arrowheads: co-localization of indicated markers.

### CRH-expressing OPCs mature into myelinating OLs

One function of OPCs in the context of brain injury is the regeneration of the OL population (Vigano *et al*., 2016). Therefore, we investigated the fate of CRH-expressing cells in more detail by evaluating their differentiation capacity using NG2/Olig2 and CC1/Olig2 double staining. At the time of their first appearance between 2 and 3 dpi, all (100 ± 0%) of tdTomato^+^/Olig2^+^ cells were also NG2^+^ (**Fig. 2f, Fig. S2c**). No co-localization with the OL marker CC1 was found. Subsequently, the proportion of NG2^+^ cells of the tdTomato^+^/Olig2^+^ population steadily decreased, reaching a minimum at 128 dpi (7.67 ± 4.10%), while the percentage of CC1^+^ cells among tdTomato^+^/Olig2^+^ cells continuously increased until 128 dpi (76.33 ± 3.67%) (**Fig. 2f,g, Fig. S2d**). CRH^+^ OLCs possessed OL-like morphologies and were present in the midbrain as well as in the white matter (WM) (**Fig. 2h**). The myelinating character of these cells was further confirmed by co-localization with the myelin proteins CNPase and MBP within their processes at 4 months post injury (**Fig. 2i**). These results indicate that CRH-expressing OPCs predominantly mature into myelinating OLs after acute injury and are highly stable once integrated. To further substantiate these findings on a single cell level, we used *CRH-Cre::Ai9* and *CRH-FlpO::NG2-CreERT2::Ai65* mice for repeated *in vivo* two-photon imaging. For cortical imaging, a cranial window in combination with an acute stab wound injury was used in *CRH-Cre::Ai9* mice **(Fig. 3a,b, Fig. S3a)**. To image WM processes, a cannula was implanted into the cortex, resembling an injury itself (**Fig. 3 a**). Using these methods, single cells were identified and followed over several weeks confirming proliferation (**Fig. S3b**), movement towards the injury site (**Fig. S3c**) and the subsequent maturation, as can be inferred from the morphological changes leading to the characteristic appearance of a myelinating OL (**Fig. 3c**). On top of that, long-term imaging in the WM of *CRH-FlpO::NG2-CreERT2::Ai65* mice (cannula implantation) revealed that mature OLs persisted over the entire imaging period (**Fig. S3d-g’’, h-k**). These findings collectively identify CRH-expressing OPCs as a highly proliferative subpopulation of OPCs, highlighting their strong propensity for oligodendrogenesis and their long-term stability following successful integration.

**Fig. 3:**
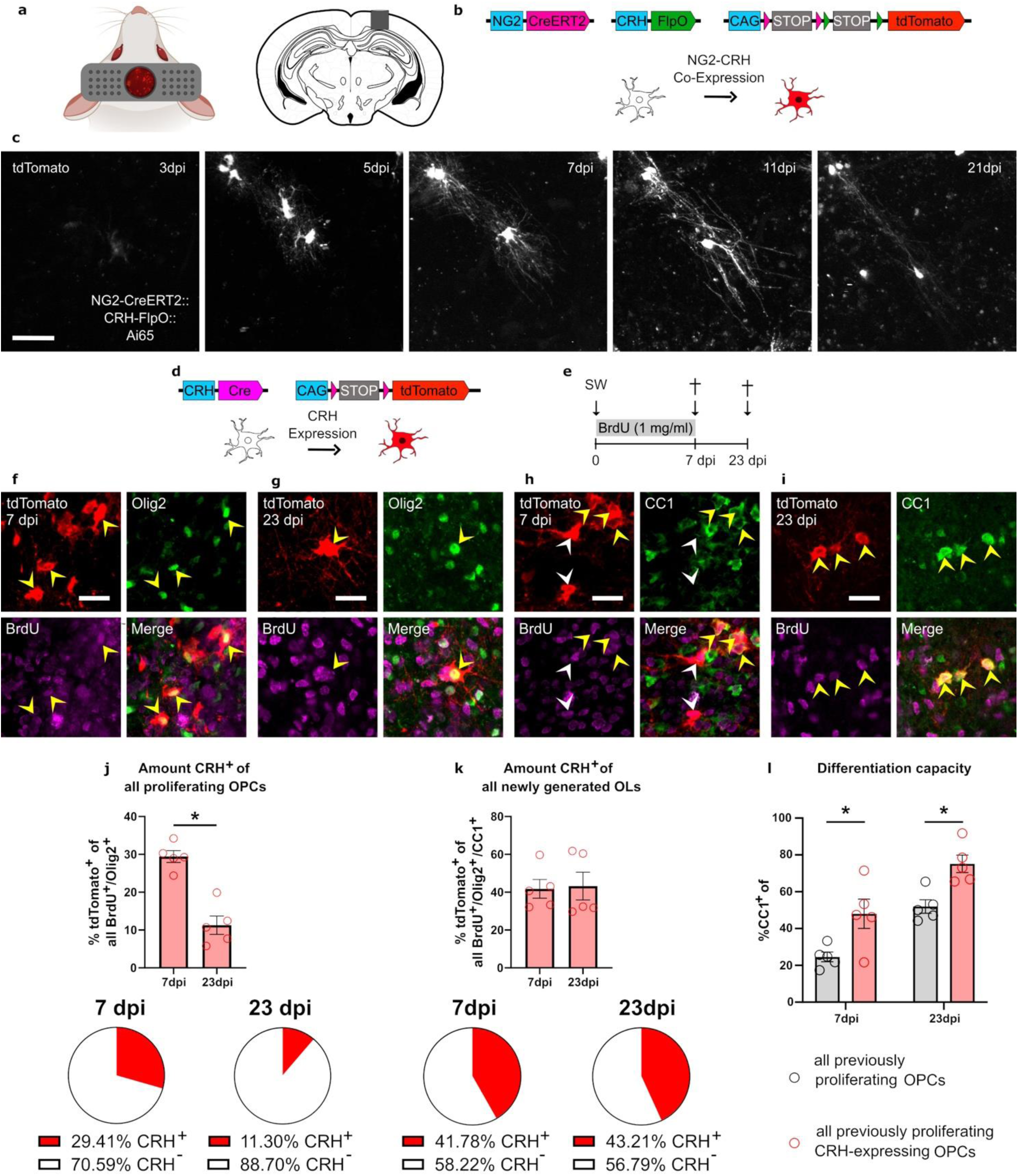
CRH^+^ OPCs show elevated differentiation capacity compared to CRH^-^ OPCs. **a**, Graphical illustration of field of view during *in vivo* 2-photon imaging (left) and the position of the imaging cannula (right). **b**, Intersectional reporter mouse line *CRH-FlpO::NG2-CreERT2::Ai65.* **c**, Representative 2-photon images of tdTomato^+^ OPCs in one ROI after implantation of hippocampal cannula between 3 and 21 dpi maturing into OLs. Scale bar, 50 µm. **d**, Graphical illustration of the *CRH Cre::Ai9* reporter mouse model. **e**, Experimental setup of label retaining experiment, **f-g**, Confocal images of tdTomato, Olig2 and BrdU colocalization at 7 (**f**) and 23 dpi (**g**). Scale bar, 20 µm. **h-i**, Confocal images of tdTomato, CC1 and BrdU colocalization at 7 (**h**) and 23 dpi (**i**). Scale bar, 20 µm. **j**, Amount CRH^+^ (tdTomato^+^) cells of all BrdU^+^/Olig2^+^ cells showing a significant decrease from 7 to 23 dpi (Welch-corrected two-tailed t-test: t_(6.9)_ = 6.26, p = 0.0005) and pie-charts illustrating proportions of CRH^+^ and CRH^-^ cells in whole population of previously proliferating OLCs over time. **k**, Amount of CRH^+^ (tdTomato^+^) cells in BrdU^+^/CC1^+^ newly generated OLs showing no change between 7 and 23 dpi and pie-charts illustrating proportions of CRH^+^ and CRH^-^ percentages in whole population of newly generated OLs. **l**, Amount of CC1^+^ cells of CRH-expressing and all previously proliferating OLCs showing a significantly elevated differentiation probability in CRH^+^ OLCs at 7 and 23 dpi (Two-way ANOVA: condition, F_(1, 16)_ = 20.29, p = 0.0004, n_TP_ = 5 mice). For all images, White arrowheads indicate cells or structures. Yellow arrowheads indicate co-localization of indicated markers. For all Two-way ANOVAs, Sidak’s post hoc test, *p < 0.05, ****p < 0.0001.

### CRH-expressing OPCs contribute substantially to OL generation following injury

To assess the size and dynamics of the CRH-expressing OLC subpopulation in relation to the entire OLC population, we utilized the robust proliferative response of OPCs following injury. We conducted a label-retaining experiment in *CRH-Cre::Ai9* mice post-insult using the intercalating substance 5-Bromo-2’-deoxyuridine (BrdU) which labels all proliferating cells (**Fig. 3d,e**). BrdU^+^ OLCs were identified at 7 and 23 dpi (**Fig. 3f-i**). The quantification of all tdTomato^+^/Olig2^+^/BrdU^+^ cells and the amount of BrdU^+^ cells in the total tdTomato^+^/Olig2^+^ cell population around the injury site revealed a decrease between 7 (174.0 ± 14.65 cells/mm^2^) and 23 dpi (80.11 ± 15.52 cells/mm^2^) and confirmed, the previously found (**Fig. 2 d,e**), high percentage (7 dpi: 94.23 ± 1.83%; 23 dpi: 93.50 ± 3.24%) of proliferating cells in the CRH-expressing OPC population (**Fig. S4a,b**). The fact that the percentage of BrdU^+^ cells in the total previously proliferating OPC population (Olig2^+^/BrdU^+^) was 29.41 ± 1.57% at 7 dpi (**Fig. 3j**) shows the significant contribution of the CRH-expressing population to all proliferating OPCs.

Furthermore, when examining the proportion of CRH-expressing OLCs in the newly generated OL population (BrdU^+^/CC1^+^), we found that they consistently accounted for approximately 40% of newly generated OLs at both 7 dpi (41.78 ± 4.94%) and 23 dpi (43.21 ± 7.33%) (**Fig. 3k**). To determine if this strong and persistent contribution was caused by a higher differentiation rate, we evaluated the percentage of CC1^+^ cells within the population of CRH-expressing and all formerly proliferating OLCs. CRH-expressing OLCs demonstrated a significantly increased likelihood of being CC1^+^ at both 7 (all: 24.57 ± 2.62 %; CRH^+^: 48.00 ± 8.00 %) and 23 dpi (all: 51.95 ± 3.68 %; CRH^+^: 75.10 ± 4.76 %) (**Fig. 3l**).

In summary, these results indicate that CRH-expressing OLCs constitute 30% of all proliferating OLCs at 7 dpi and, due to their higher differentiation capacity, contribute approximately 40% to the population of newly generated OLs following injury.

### Induction of CRH expression in OPCs is an early response to brain injury

After defining the key properties of CRH-expressing OPCs with regards to proliferation and maturation, we focused on the injury-induced expression of the neuropeptide CRH itself. To this end, we took advantage of the *CRH-Venus* mouse line (Kono et al., 2017) in which, Venus was inserted in the *Crh* locus, which facilitates direct monitoring of CRH expression via the reporter (**Fig. 4a, Table S1**). This enabled us to visualize the distribution of CRH^+^ cells and provided a proxy to assess CRH expression kinetics as these are directly correlated with the Venus expression driven by the endogenous *Crh* promoter. Following injury, Venus^+^/NG2^+^ cells were identified around the injury site (**Fig. 4a**). Furthermore, CRH was identified in Venus^+^ cells, validating Venus as a proxy for CRH expression (**Fig. 4b,b’**). To assess CRH expression kinetics, Venus^+^/Olig2^+^ OPCs (at early timepoints N_CRH_^+^ _Olig2_^+^ = N_CRH_^+^_NG2_^+^, **Fig. 2f**) were quantified around the injury site at different timepoints between 12 and 168 hours post injury (hpi) (**Fig. 4c**). The first Venus^+^ OPCs were detected as early as 12 hpi, indicating that CRH is expressed as an immediate reaction to the injury (**Fig. 4c**). Between 12 (12 ± 2.08 cells) and 48 hpi (80.33 ± 12.39 cells), the number of Venus-expressing OPCs increased. Comparison to a recent sequencing study analyzing early timepoints following spinal cord injury confirmed an early upregulation of CRH (Li et al., 2022). After 96 (6.67 ± 0.88 cells) and 168 hpi (4.67 ± 1.20 cells) only few cells were traceable, suggesting that Venus and, thus, CRH expression predominantly occurs within the first 3 to 4 dpi. Notably, at 12 and 24hpi all cells were present as individual single cells, whereas at 36 (46.25 ± 0.94%) and 60 hpi (76.74 ± 3.22%), a high proportion of cells were part of a cluster, likely emerging from proliferation (**Fig. S4c**). Indeed, immuno staining revealed that a large proportion of Venus^+^ cells were also Ki67^+^ but only starting from 36 hpi (39.73 ± 2.83%) onwards (**Fig. S4d**). These results, in combination with the observation that the vast majority of Olig2^+^/tdTomato^+^ cells was Ki67^+^ at 2 dpi (98 ± 1.75%) in the *CRH-Cre::Ai9* model (**Fig. 2e**), indicate that in the sequence of reactions to injury CRH expression precedes proliferation.

**Fig. 4:**
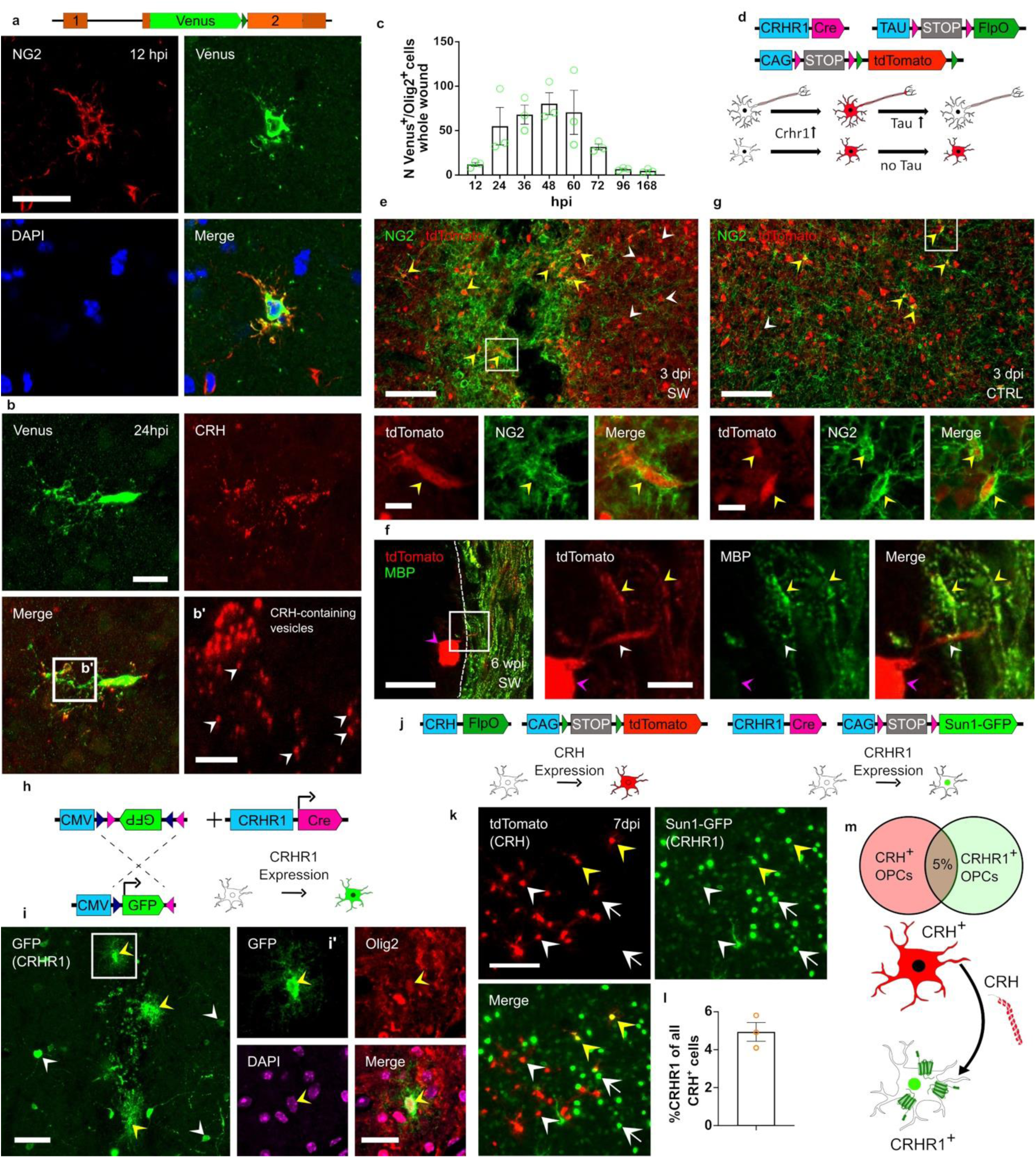
OPC-derived CRH targets distinct CRHR1^+^ OPC population. **a**, *CRH-Venus* reporter mouse model. Representative confocal images of NG2/GFP co-localization at 12 hpi. Scale bar, 20 µm. **b**, Confocal image of GFP-expressing cell at injury site in *CRH-Venus* mice stained for CRH protein. Scale bars, 20 µm (overview), 5 µm (close up). **c**, Quantification of GFP^+^/Olig2^+^ cells around the injury site between 12 and 168 hpi (One-way ANOVA: F_(7, 16)_ = 5.765, p = 0.0018). n_TP_ = 3 mice. **d**, *CRHR1-Cre::Tau-LSL-FlpO::Ai9* mouse model. **e**, Confocal image of NG2 staining around the wound identifying cells as CRHR1-expressing OPCs, Scale bars: 100 µm (overviews), 10 µm (insets). **f**, Confocal image of colocalization of MBP^+^ and tdTomato^+^ process at injury site at 23 dpi following tissue expansion. Scale bar, 100 µm. **g**, Confocal image of NG2^+^/tdTomato^+^ cells at the uninjured contralateral side at 3 dpi identifying cells as CRHR1-expressing OPCs. Scale bars: 100 µm (overviews), 10 µm (insets). **h**, Injected AAV construct and *CRHR1-Cre* mouse line for the confirmation of CRHR1 expression in OPCs. **i**, Overview of AAV injection site. Yellow arrow heads: GFP-(CRHR1^+^) expressing non-neuronal cells. White arrow heads: GFP-(CRHR1^+^) expressing neuronal cells. Scale bar, 50 µm. **i’**, Confocal image of GFP^+^/Olig2^+^ OLC at injury site confirming co-localization of markers. Scale bar, 20 µm. **j**, *CRH-FlpO::CRHR1-Cre::Ai65F::Sun1-GFP* reporter mouse model to study the percentage of overlap between the CRH- and CRHR1-expressing populations of OLCs. **k**, Confocal images of injury site in *CRH-FlpO::CRHR1-Cre::Ai65F::Sun1-GFP* mice at 7 dpi showing expression and co-expression of tdTomato (CRH) and Sun1-GFP (CRHR1). Scale bar, 100 µm. Arrowheads show tdTomato^+^/Sun1-GFP^-^ (white) and tdTomato^+^/Sun1-GFP^+^ (yellow) cells. Arrows show Sun1-GFP^+^/tdTomato^-^ cells. **l**, Quantification of CRHR1^+^/CRH^+^ of all CRH^+^ cells. N = 3 animals. **m**, Population overlap between CRH- and CRHR1-expressing OPCs and potential directionality of interaction. White arrowheads indicate cells or structures. Yellow arrowheads indicate co-localization of indicated markers. Purple arrowhead indicates soma.

### CRHR1-expressing OPCs resemble a distinct population and are potential targets of OPC-derived CRH

The canonical high-affinity receptor of CRH is the CRHR1. To identify potential target cells of CRH-release we investigated the presence of CRHR1-expressing glial cells in vicinity of the wound. Therefore, we used the *CRHR1-Cre::Tau-LSL-FlpO::Ai9* mouse model, in which tdTomato expression is activated in all CRHR1^+^ cells, but selectively deleted from CRHR1^+^ neurons. (**Fig. 4d, Table S1**). When analyzing stab wounds at 3 dpi, we found CRHR1-expressing cells aggregating around the injury site that were identified as OPCs by NG2 staining (**Fig. 4e**). These CRHR1-expressing OPCs were also shown to differentiate into myelinating OLs at 6 wpi by MBP staining following tissue expansion (Chen et al., 2015; Delling et al., 2024) **(Fig. 4f)**. To our surprise, CRHR1^+^/NG2^+^ cells were also present on the non-injured contralateral site and also throughout the brain under non-injury conditions (**Fig. 4g**). To better understand this potential interaction between OPC-derived CRH and CRHR1 on OPCs we tested whether CRHR1-expression persisted in adulthood. Due to the lack of reliable CRHR1-specific antibodies, we had to choose other approaches for detection of CRHR1 expression (Refojo et al., 2011). Thus, we validated the expression of CRHR1 in OLCs by: i) AAV-mediated reporting of CRHR1 expression and ii) direct visualization of CRHR1 expression in *CRHR1^ΔEGFP^* mice. For AAV-mediated reporting we used an AAV harboring the DIO-GFP system, which was injected in *CRHR1-Cre* mice (**Fig. 4h**). Following Cre-mediated recombination, GFP was activated in all CRHR1-expressing cells.. CRHR1^+^ OLCs were specifically identified by Olig2 colocalization (**Fig. 4i,i’**). To assess CRHR1 expression in OPCs under physiological conditions we took advantage of the *CRHR1^ΔEGFP^* mouse line which is characterized by a knock-in of GFP into the *Crhr1* locus, which serves as valid proxy for CRHR1 expression **(Fig. S4e, Table S1)** (Refojo *et al*., 2011). Co-staining with PDGFRα showed clear co-localization with GFP-expressing cells of the WM, identifying these cells as OPCs (**Fig. S4f,f’**).

The identification of CRHR1-expressing OPCs surrounding the injury site raised the question whether ligand and receptor are cellularly co-expressed or exist in distinct OPC populations. To quantify the overlap, we used the *CRH-FlpO::CRHR1-Cre::Ai65F::Sun1-GFP* mouse line in which CRH expression is reported by tdTomato and CRHR1 expression by GFP fused to the nuclear membrane localization marker SUN1 (**Fig. 4j, Table S1**). TdTomato^+^ (CRH) and GFP^+^ (CRHR1) cells were identified around the injury site, but only showed an overlap of 4.9 ± 0.49% (**Fig. 4k,l**), thereby clearly separating the CRH- and CRHR1-expressing populations of OPCs (**Fig. 4m**). Taken together, these observations evidently demonstrate that CRHR1 is expressed in a distinct OPC population, qualifying it as potential target of CRH released following injury.

### CRHR1^+^ OPCs show a delayed differentiation following injury

To investigate the differences between CRHR1- and CRH-expressing OPCs following injury, we conducted a label-retaining experiment using *CRHR1-Cre::Sun1-GFP* animals (**Fig. 5a,b; Table S1**). These animals were subjected to injury, treated with BrdU for 3 or 7 days and sacrificed at 3, 7, and 23 dpi. GFP^+^/BrdU^+^/Olig2^+^ and GFP^+^/BrdU^+^/CC1^+^ cells were identified at all timepoints (**Fig. 5c-f**).

**Fig. 5:**
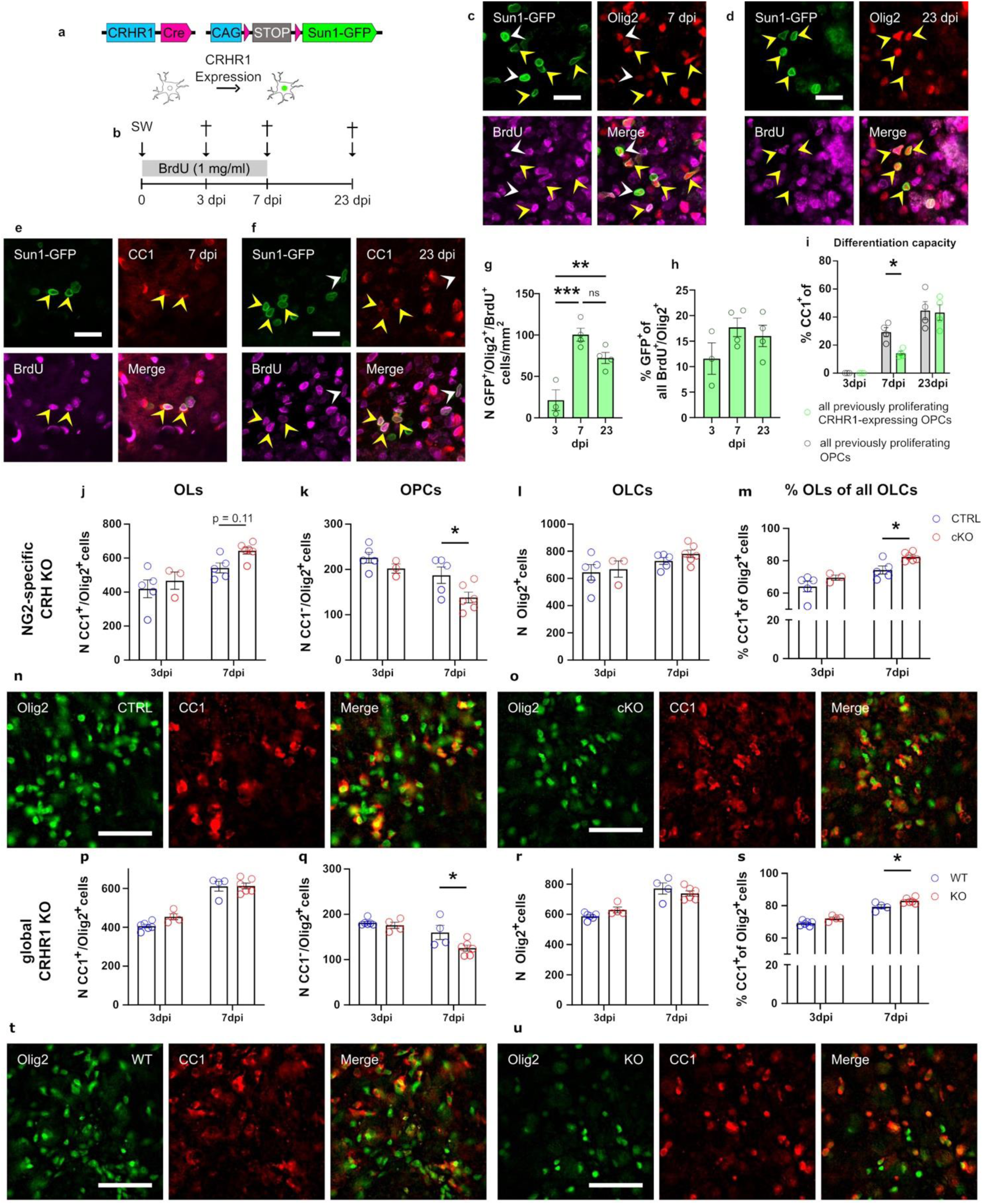
Injury-related population dynamics of CRHR1^+^ OLCs and effect of CRH/CRHR1 modulation. **a**, *CRHR1-Cre::Sun1-GFP* mouse model. **b**, Experimental timeline of BrdU label retaining experiment. **c** and **d**, Confocal images of Sun1-GFP^+^ (CRHR1^+^)/Olig2^+^/BrdU^+^ cells at 7 (**c**) and 23 dpi (**d**). **e** and **f**, Confocal images of Sun1-GFP^+^ (CRHR1^+^)/CC1^+^/BrdU^+^ cells at 7 (**e**) and 23 dpi (**f**). **g**, Quantification of BrdU^+^ CRHR1-expressing OLCs at 3, 7 and 23 dpi (One-way ANOVA: F_(2, 8)_ = 19.6, p = 0.0008), **h**, Amount of CRHR1 (GFP)-expressing OPCs of all proliferating (BrdU^+^/Olig2^+^) OPCs., Amount of CC1^+^ cells of CRHR1-expressing and all previously proliferating OLCs (Two-way ANOVA: condition, F_(1, 16)_ = 2.6, p = 0.1295, n_TP_ = 4 mice). **j**, Number of CC1^+^/Olig2^+^ cells at 3 and 7 dpi in NG2-specific CRH cKO vs CTRL animals (Two-way ANOVA: genotype, F_(1,15)_= 3.76, p = 0.0714, n_CTRL 3dpi_ = 5, n_cKO 3dpi_ = 5, n_CTRL 7dpi_ = 5, n_cKO 7dpi_ = 6). **k**, Number of CC1^-^/Olig2^+^ cells in NG2-specific CRH cKO compared to CTRL animals (Two-way ANOVA: genotype, F_(1,15)_= 6.3, p = 0.024). **l**, The total number of Olig2^+^ cells in cKO and CTRL animals. **m**, Percentage of CC1^+^ of all Olig2^+^ cells in NG2-specific CRH cKO and CTRL animals (Two-way ANOVA: genotype, F_(1,15)_= 8.078, p = 0.0124). **n and o,** Representative images of Olig2/CC1 staining in CTRL (**e**) and cKO (**f**) animals. Scale bars, 50 µm. **p**, Number of CC1^+^/Olig2^+^ in global CRHR1 KO and WT animals (n_WT 3dpi_ = 6, n_KO 3dpi_ = 4, n_WT 7dpi_ = 4, n_KO 7dpi_ = 6). **q**, Number of CC1^-^/Olig2^+^ cells in global CRHR1 KO animals (Two-way ANOVA: genotype, F_(1,16)_= 6.244, p = 0.0237). **r**, Number of Olig2^+^ in CRHR1 KO and WT animals. **s**, % of CC1^+^ cells of Olig2^+^ cells in global CRHR1 KO and WT animals (Two-way ANOVA: genotype, F_(1,16)_= 16.08, p = 0.001). **t and u**, Representative images of CC1/Olig2 staining in WT (**t**) and KO (**u**) animals. Scale bars, 50 µm. For all Two-way ANOVAs, Sidak’s post hoc test, *p < 0.05, ****p < 0.0001. For all images, White arrowheads indicate cells or structures. Yellow arrowheads indicate co-localization of indicated markers. For all Two-way ANOVAs, Sidak’s post hoc test, *p < 0.05, ****p < 0.0001.

First, we assessed the number of previously proliferating CRHR1^+^ OLCs (GFP^+^/Olig2^+^/BrdU^+^) noting a significant increase in cell numbers from 3 dpi (21.25 ± 12.53 /mm^2^) to 7 dpi (100.4 ± 7.80 /mm^2^), followed by a non-significant decrease towards 23 dpi (72.31 ± 6.70 /mm^2^; **Fig. 5g**).

We then analyzed the proportion of CRHR1-expressing OLCs (GFP^+^/BrdU^+^/Olig2^+^) within the population of all previously proliferating OLCs (BrdU^+^/Olig2^+^). The proportion increased slightly from 11.57 ± 5.34% at 3 dpi to 17.70 ± 3.62% at 7 dpi, subsequently remaining steady until 23 dpi (16.01 ± 2.09%; **Fig. 5h**) showing that the CRHR1-expressing population of OPCs contributed significantly to the population of proliferating OPCs.

To evaluate the differentiation capacity of CRHR1-expressing OPCs, we compared the percentage of CC1^+^ cells in the overall BrdU^+^ and within the CRHR1^+^ population. Our analysis revealed a significantly lower percentage of CC1^+^ cells among CRHR1-expressing OLCs at 7 dpi (All: 29.27 ± 3.22%; CRHR1^+^: 14.18 ± 1.43%), a difference that dissipated by 23 dpi (All: 44.60 ± 6.42%; CRHR1^+^: 43.25 ± 5.57%; **Fig. 5i**). This temporary disparity in the probability of a cell being CC1^+^ between the two populations suggests a delay in the differentiation of CRHR1^+^ OPCs, rather than a fundamental difference in their differentiation potential following injury.

### Disruption of the CRH/CRHR1 system affects the OPC population around the injury site

After the identification and characterization of CRHR1-expressing OPCs around the injury site as potential targets of OPC-derived CRH, we investigated the functional impact of the inhibition of the CRH/CRHR1 system on OLCs following insult.

To this end, we pursued different loss-of-function (LOF) approaches using i) a conditional tamoxifen-inducible NG2-specific CRH KO (*CRH ^NG2-cKO^*) and ii) a global constitutive CRHR1 KO (*CRHR1^ΔEGFP^*) (**Table S1**). Before assessing the effect of and the NG2-specific CRH KO, the recombination efficiency following tamoxifen treatment was assessed and shown to be at 66.3 ± 1.6 % (**Fig. S5a,b**). First, we compared the number of OLs, OPCs, OLCs and the amount of OLs in the whole population of OLCs (**Fig. 5j-m**). In *CRH ^NG2-cKO^* mice, no significant difference was identified in OL or OLC numbers, but the number of OPCs around the injury site was significantly reduced in cKO animals at 7 dpi **(Fig. 5k,n,o**). This lower number of OPCs was accompanied by significantly increased proportion of OLs of all OLCs in cKO animals at 7 dpi (**Fig. 5m**).

Similar to *CRH ^NG2-cKO^* mice, the global loss of CRHR1 expression in *CRHR1^ΔEGFP^* led to a significant reduction in OPCs and also to a higher proportion of OLs of all OLCs at 7 dpi (**Fig. 5q,t,u**), while the total number of OLs and OLCs was unchanged (**Fig. 5p,r**). None of these effects were observed on the non-injured, contralateral side (**Fig. S5c-f**). These results indicated that a decreased CRH/CRHR1 signaling influences the OLC population following injury by altering the dynamics of OPCs.

### CRH/CRHR1 system inhibition increases the generation and survival of newly formed OLs in an injury-size dependent manner

The observed reduction of OPCs following CRH/CRHR1 system blockade could have been caused by a reduced proliferation between 3 and 7 dpi or by a faster differentiation rate of OPCs into OLs. To elucidate the cause, we performed a label retaining experiment using *CRHR1^ΔEGFP^* KO mice (**Fig. 6a**). Neither the number of BrdU^+^, all BrdU^+^/Olig2^+^ or BrdU^+^/Olig2^+^/CC1^-^ showed any significant difference between WT and KO animals. suggesting equal proliferative activation of OPCs (**Fig. 6b, Fig. S6a,b**). We then quantified the number of BrdU^+^/Olig2^+^/CC1^+^ cells around the injury site and found a significant increase in these newly generated OLs at 7 dpi which was absent at 6 wpi (**Fig. 6c**). When limiting the analysis (**Fig. 6e**), to the inner 150 µm of the wound, a significant increase in the total number of OLs (CC1^+^/Olig2^+^ cells) was present in KO animals (**Fig. 6e**). The mean difference in the total number of OLs was twice as high as the difference in newly generated ones (BrdU^+^/Olig2^+^/CC1^+^ cells) suggesting that direct differentiation from OPCs without proliferation and concomitant BrdU integration also contributed to these differences. When analyzing the number of newly generated OLs in the first 150 µm, we found a significantly higher number in KO animals compared to WT animals (**Fig. 6f**). Overall, the observed reduction of OPCs in the CRH as well as CRHR1 KO animals were most likely caused by an increased differentiation velocity at early post-injury stages.

**Fig. 6:**
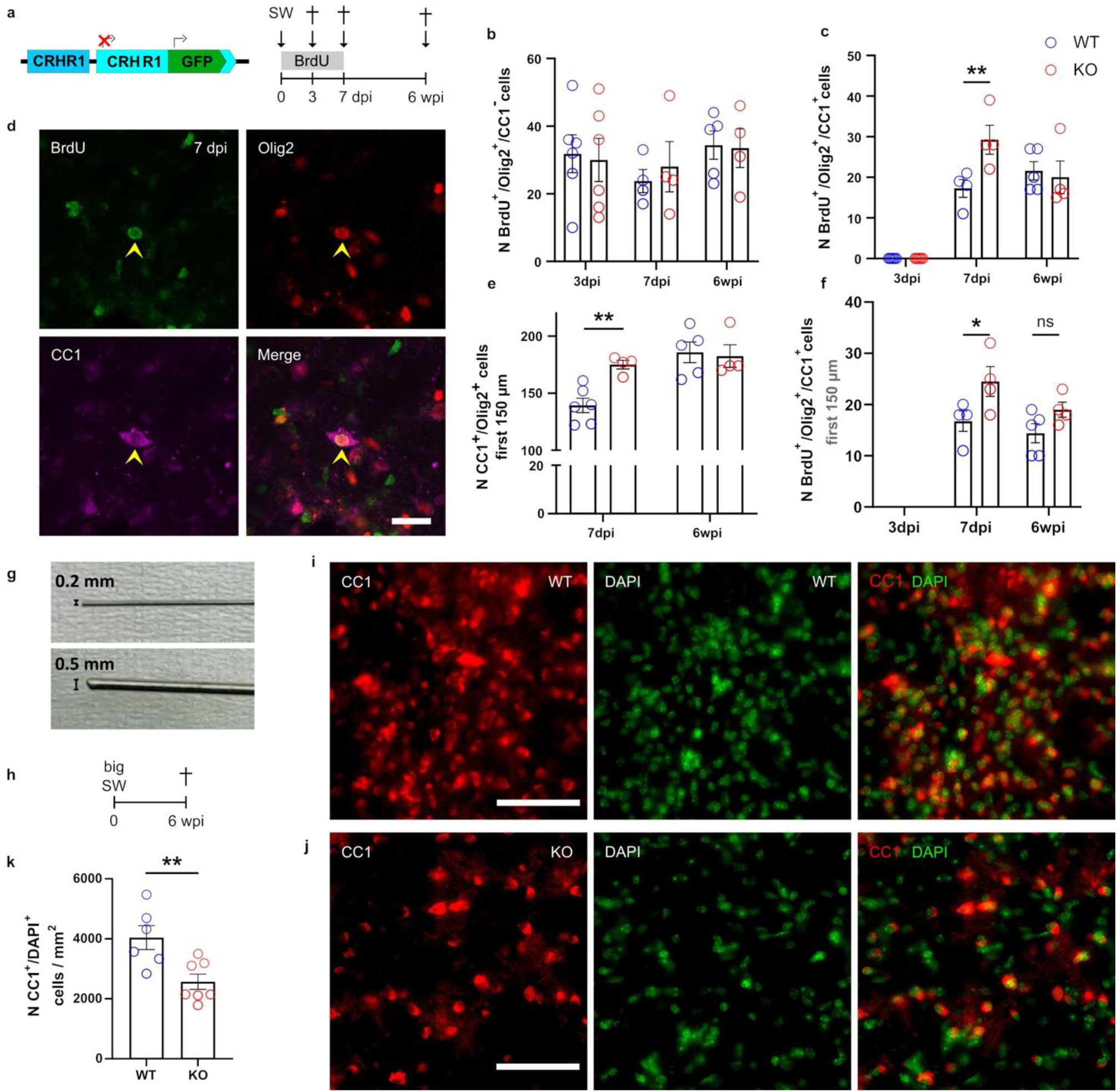
CRH/CRHR1 system regulates early differentiation and long term-survival after injury. **a,** *CRHR1^ΔEGFP^* mouse model and experimental scheme. **b**, Quantification of BrdU^+^/Olig2^+^/CC1^-^ cells in 300 µm radius around injury site. **c**, Quantification of BrdU^+^/Olig2^+^/CC1^+^ cells around injury site (Two-way ANOVA: time, F_(2,23)_ = 77.67, p < 0.003; genotype, F_(2,23)_ = 3.841, p = 0.0622, n_WT 3dpi_ = 6, n_KO 3dpi_ = 6, n_WT 7dpi_ = 4, n_KO 7dpi_ = 4, n_WT 6wpi_ = 5, n_KO 6wpi_ = 4). **d**, Confocal image of BrdU/Olig2/CC1 staining at 7 dpi. **e-f**, Quantification of CC1^+^/Olig2^+^ cells in whole wound area (**e**) and within 150 µm (**f**) (Two-way ANOVA: time, F_(1,15)_ = 12, p = 0.0035; genotype, F_(1,15)_ = 4.316, p = 0.0553, n_WT 7 dpi_ = 6, n_KO 7 dpi_ = 4, n_WT 6wpi_ = 5, n_KO 6wpi_ = 4). **g**, Binocular image of small (0.2 mm) and big (0.5 mm) Hamilton syringe. **h**, Experimental setup to test influence of 2.5×increased injury size. **i** and **j**, Confocal images of CC1^+^/DAPI^+^ cells around the injury site in WT (**i**) and KO animals (**j**). Scale bars, 50 µm. **k**, Quantification of CC1^+^/DAPI^+^ cells/mm^2^ in the first 50 µm around the injury site at 6 wpi (Two-tailed Students t-test: t_(11)_ = 3.21, p = 0.0083). Yellow arrowheads indicate co-localization of indicated markers. For all Two-way ANOVAs, Sidak’s post hoc test, *p < 0.05, ****p < 0.0001.

To assess the influence of the injury size on the long-term survival of OLs, we performed another experiment in *CRHR1^ΔEGFP^* animals using a 2.5× bigger cannula (diameter 0.5 mm) (**Fig. 6g,h**). Animals were sacrificed at 6 wpi and the number of CC1^+^ OLs was assessed on coronal sections to increase the visible part of the injury. The analysis of the injury site showed a significant decrease in KO compared to WT animals (**Fig. 6i-k**). Thus, in CRHR1 KO animals, the long-term survival of newly generated OLs is reduced in a threshold-dependent manner, depending on the injury size and the corresponding demand for remyelination. This decrease was likely caused by the CRHR1 KO-induced premature differentiation of OPCs into OLs observed in the previous experiment (**Fig. 6c-f**).

### CRHR1-expressing OLCs increase their numbers in the whole OLC population under non-injury conditions

Because CRH-expressing OPCs are only present in the murine brain following acute injury, but CRHR1 expressing OPCs as well as other sources of CRH, e.g. neurons, are present under naive conditions, we next sought to investigate their functional role under non-injury conditions. To this end we quantified the number of CRHR1-expressing OLCs at different postnatal timepoints (1.5, 3 and 5 months) in *CRHR1-Cre::Tau-LSL-FlpO::Ai9* mice (**Fig. 7a**). CRHR1-expressing OLCs were found at all ages (**Fig. 7b-f**). Firstly, the total number of CRHR1-expressing OPCs was quantified at all timepoints in the cortex (CX), WM including corpus callosum, capsule and anterior commissure as well as in the MB. The number of CRHR1-expressing OPCs increased over time, showing a significant gain between 3 and 5 months in the CX (2.61 ± 0.96 to 19.67 ± 3.54 cells/ mm^2^) and MB (9.78 ± 2.13 to 28.44 ± 8.30 cells/ mm^2^) (**Fig. S7a**). This resulted in a significant increase in the percentage of tdTomato^+^ OPCs of all OPCs in the CX (1.88 ± 0.35 to 7.07 ± 1.44%) and MB (4.36 ± 0.87 to 10.42 ± 2.64%) but not in the WM (2.09 ± 0.89% to 4.81 ± 0.39 %) between 1.5 and 5 months (**Fig. S7b**). In addition, we quantified the total number of tdTomato^+^/Olig2^+^ cells, which differed between distinct brain regions and ages. In the CX, the cell numbers were lowest at all timepoints, while the numbers in WM and MB were comparable at 1.5 and 3 months. Generally, the number of tdTomato^+^/Olig2^+^ cells significantly increased towards 5 months of age, which was more pronounced in the WM (100.70 ± 37.35 to 362.78 ± 95.95 cells/mm^2^) than in the MB (124.06 ± 19.62 to 217.5 ± 33.17 cells/mm^2^) (**Fig. 7g**). As the number of OLs increases with age, we investigated whether the increase in CRHR1^+^ OLCs paralleled that of the overall OL population (Tripathi et al., 2017). We quantified the total number of Olig2^+^ cells (**Fig. S7c**) and calculated the percentage of CRHR1^+^/Olig2^+^ cells, which indeed significantly increased in WM and MB over time (**Fig. 7h**). This increase in the proportion of CRHR1^+^/Olig2^+^ cells suggested that CRHR1^+^ OLCs increased their number at a higher rate than the population of CRHR1^-^ OLCs, implicating that the CRHR1^+^ OPC subpopulation may play an important role in adult OL maturation and potentially also myelination.

**Fig. 7:**
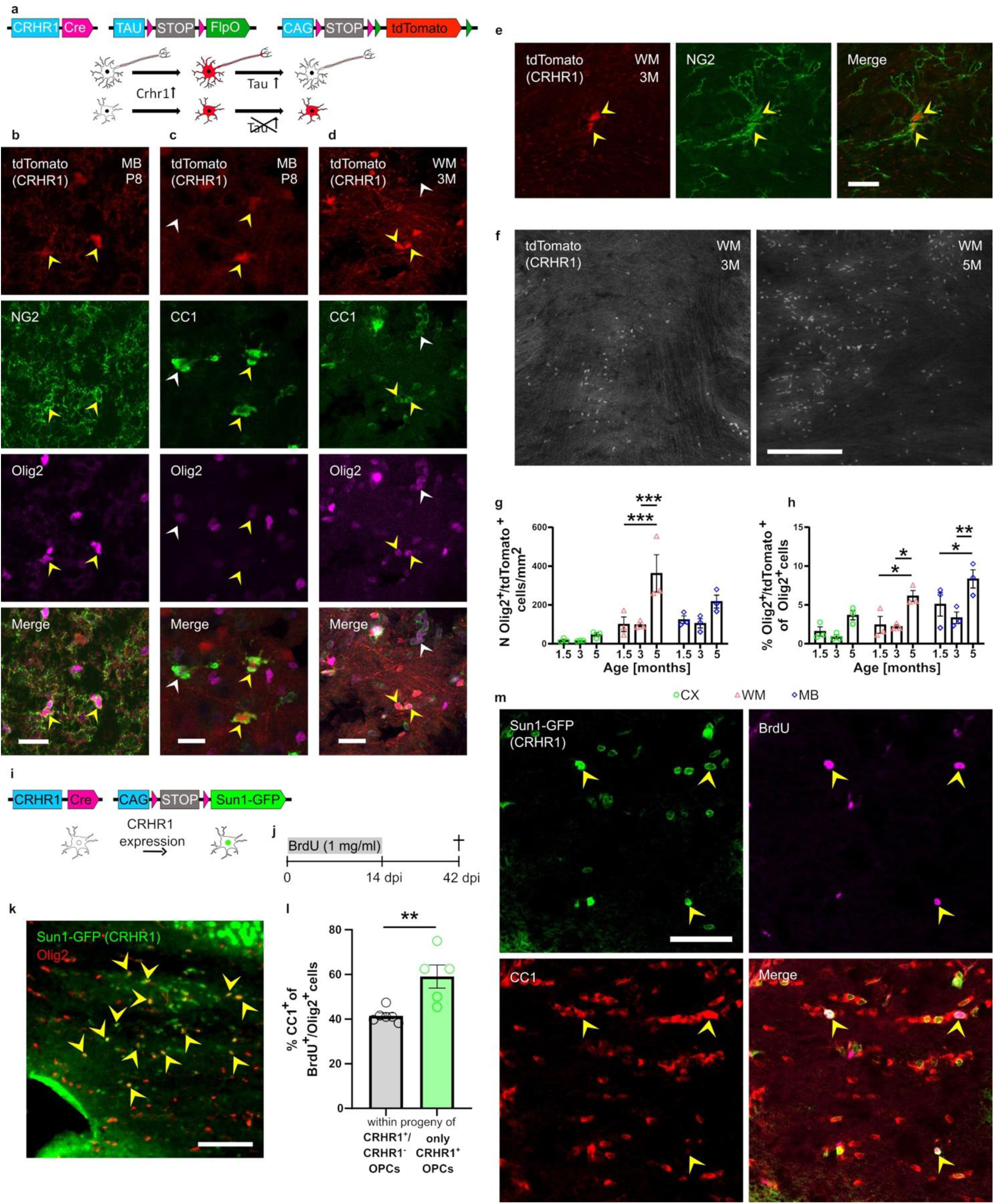
Population dynamics of CRHR1^+^ OLCs under non-injury conditions. **a,** *CRHR1::Tau-LSL-FlpO::Ai9* reporter mouse model. **b-e**, Confocal images showing co-localization of tdTomato (CRHR1) with NG2 and CC1 at P8 and at 3 months. Scale bar, 20 µm. **f**, Maximum intensity projections of confocal images of WM at 3 and 5 months of age. Scale bar, 200 µm. **g**, Quantification of NG2^+^/tdTomato^+^ cells at 1.5, 3 and 5 months of age in CX, WM and MB (Two-way ANOVA: time, F_(2, 18)_ = 10.34, p = 0.001, n_TP_ = 3 mice). **h**, % Olig2^+^/tdTomato^+^ of all Olig2^+^ cells in WM and MB (Two-way ANOVA: time, F_(2, 18)_ = 16.65, p < 0.0001; region, F_(2, 18)_ = 12.54, p = 0.0004, n_TP_ = 3 mice). **i**, *CRHR1-Cre:Sun1-GFP* reporter mouse model. **j,** Label retaining experiment using BrdU in *CRHR1-Cre:Sun1-GFP* to study OPC differentiation under non-injury conditions**. k,** Confocal images showing aggregation of Sun1-GFP^+^ (CRHR1^+^)/Olig2^+^ cells in CC. Scale bar, 50 µm. **l**, % CC1^+^ of BrdU^+^/Olig2^+^ and all BrdU^+^/Sun1-GFP^+^ (CRHR1^+^) at 42 dpi, (Two-tailed unpaired t-test: t_(9)_ = 3.569, p = 0.0060, N_animals_ = 5). **m**, Confocal images of CC1^+^/BrdU^+^/Sun1-GFP^+^ (CRHR1^+^) cells in CC. Scale bar, 50 µm. For all images, White arrowheads indicate cells or structures. Yellow arrowheads indicate co-localization of indicated markers. For all Two-way ANOVAs, Sidak’s post hoc test, *p < 0.05, ****p < 0.0001.

### Increased number of CRHR1-expressing OLCs under non-injury conditions is caused by an elevated differentiation rate

To further investigate the cause of the observed increase in CRHR1-expressing OLCs, we conducted a label-retaining experiment in *CRHR1-Cre::Sun1-GFP* mice by treating them with BrdU for two weeks, followed by a retaining phase of 4 weeks (**Fig. 7i,j**). We then compared the differentiation rate of CRHR1-expressing OPCs to that of all OPCs in the CC (**Fig. 7k**). Assessment of the percentage of CC1^+^ cells in the population of formerly proliferating cells (OPCs) revealed a significantly higher percentage of differentiated cells in the CRHR1-expressing population (CRHR1^+^: 59.09 ± 5.22%, All: 41.53 ± 1.28%) after the 4-week retaining period (**Fig. 7l,m**). Combined with previous results following acute injury, this suggests that CRHR1 may act as an inhibitory regulator, preventing premature oligodendrogenesis under certain conditions, which is critical for adequate differentiation and thus long-term survival.

### Inhibition of CRH/CRHR1 interaction under non-injury conditions leads to premature myelination and long-lasting changes in the myelin structure

While CRH-expression in OPCs can only be observed following injury, neurons represent the major source of CRH under non-injury conditions. To test the effect of CRH/CRHR1 signaling in OPCs under non-injury conditions and its effect on oligodendrogenesis, we used *CRHR1^ΔEGFP^* mice. Because the main developmental oligodendrogenesis and myelination occurs during early postnatal stages (de Faria et al., 2021), we first sacrificed *CRHR1^ΔEGFP^* mice at postnatal day 8 (p8) and assessed the number of OLs in CRHR1 WT and KO animals (**Fig. 8a,b**). Intriguingly, we found increased numbers of OLs in the central corpus callosum (CC) in CRHR1 KO mice (**Fig. 8c,d**). This suggests that CRH/CRHR1 signaling inhibits premature oligodendrogenesis not only following injury but also during early postnatal development.

**Fig. 8:**
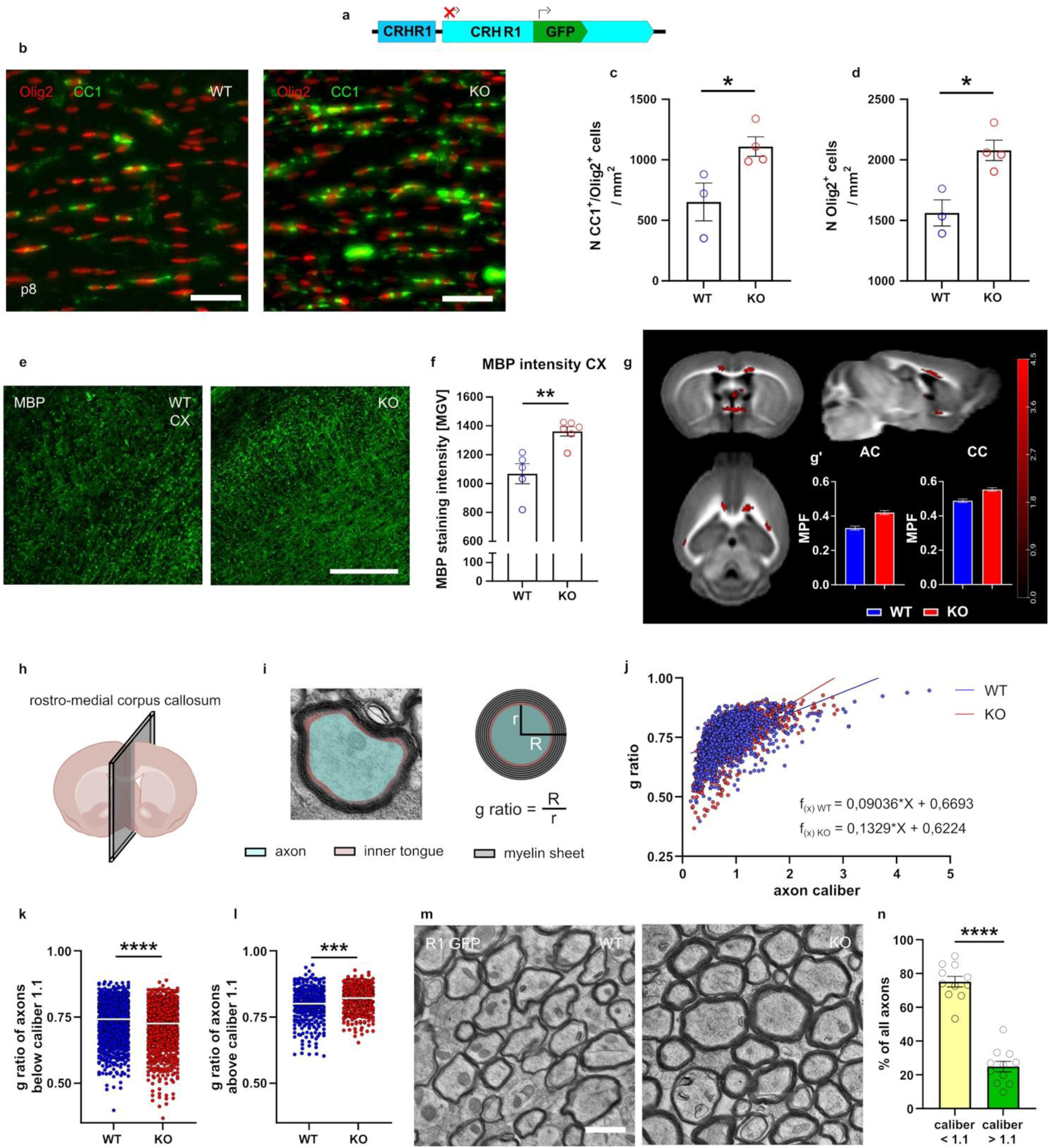
CRH/CRHR1 system regulates early postnatal and adult myelination. **a**, *CRHR1^ΔEFGP^* mouse model a *de facto* KO of CRHR1. **b**, Confocal images of CC1^+^/Olig2^+^ cells in CC of *CRHR1^ΔEGFP^* WT and KO animals at p8. Scale bar, 50 µm. **c** and **d**, Quantification of CC1^+^/Olig2^+^ (**c**) and all Olig2^+^ (**d**) cells/mm^2^ in CC (Two-tailed Students t-test: n_CC1/Olig2_: t_(5)_ = 2.82, p = 0.037; n_Olig2_: t_(5)_ = 3.85, p = 0.012). **e**, Confocal image of MBP staining in CX of WT and KO (*CRHR1^ΔEGFP^*) mice. Scale bar, 100 µm. **f**, Quantification of staining intensity of MBP in CX of WT and KO mice (Two-tailed Students t-test: t_(9)_ = 4.07, p = 0.0028; n_animals/condition_ = 5-6). **g**, Graphical illustration of calculated T-contrast (p_FDR cluster_ < 0.05, collection threshold of p < 0.01) on brain matrix. **g’**, Plotted MPF peak voxel values in AC and CC. **h**, Cutting plane for EM analysis. **i**, Illustration of location of axon, inner tongue and myelin sheet and of g ratio calculation (with R = Radius myelin sheet and r = radius axon). **j**, g ratio in dependence of axon caliber of WT and KO animals. **k** and **l**, g ratio of axon with caliber smaller (**k**) and bigger (**l**) than 1.1 µm (Mann-Whitney U test: smaller 1.1 µm: Mann-Whitney U = 423035, p < 0.0001; bigger 1.1 µm: Mann-Whitney U = 42738, p = 0.0002, n_animals/GT_ = 5-6). **m**, EM image of myelinated axons in medial CC in WT and KO animals. Scale bar, 1 µm. **n**, % of all axons with a caliber below and above 1.1 µm (Two-tailed Students t-test: t_(20)_ = 11.31, p < 0.0001; N_total_ = 11 mice).

Next, we assessed whether this CRHR1 KO-dependent premature oligodendrogenesis also impacted on adult myelination patterns. We first conducted anti MBP stainings on brain sections of adult *CRHR1^ΔEGFP^* animals and assessed OL numbers as well as potential intensity and coverage differences in CX and WM. The analyses revealed that the MBP staining intensity was significantly increased in the CX of CRHR1 KO animals (WT: 1068 ± 69.36 MGV; KO: 1362 ± 32.44 MGV) (**Fig. 8e,f**). In the WM, MBP intensity differences were not detectable, likely due to the high default background intensity (data no shown). To overcome these intensity-connected drawbacks, we used magnetic resonance imaging (MRI) to measure myelin content differences more globally. A specific MRI method is the magnetization transfer (MT), which can be used to estimate the Macromolecular Proton Fraction (MPF). MPF represents the most sensitive biomarker of myelin density in MRI and has been shown to be superior to other methods like diffusion tensor imaging derived fractional anisotropy (Hertanu et al., 2023; Khodanovich et al., 2017; Soustelle et al., 2021; Yarnykh, 2012; Yarnykh, 2016). Comparison between CRHR1 WT and KO mice revealed significantly increased myelin content in various brain regions, including the medial and lateral CC and AC (**Fig. 8g,g’**). To further understand the underlying differences in these myelin alterations and to get insight in potential structural differences in myelin, we performed electron microscopy (EM) of the rostro-medial CC of adult *CRHR1^ΔEGFP^* mice (**Fig. 8h**). Analysis of the g ratio in relation to axon caliber showed significant differences in the linear regression slopes between WT and KO animals (**Fig. 8i,j**). To better understand the differences in the g ratio in different axon calibers, we calculated the intersection point between the two regression curves. The calculated value of 1.1 µm was used to separate the axons in two populations, smaller or larger than an axon caliber of 1.1 µm. We then compared the average g ratio of axons smaller or larger than 1.1 µm between genotypes and found that smaller caliber axons in KO mice had a significantly smaller g ratio (thicker myelin) (WT: 0.73 ± 0.0023; KO: 0.71 ± 0.0026), while larger caliber axons had a significantly increased g ratio (thinner myelin) (WT: 0.80 ± 0.0036; KO: 0.82 ± 0.0028) (**Fig. 8k,l**).

To understand the impact of these g ratio changes on overall myelin content, we examined the distribution of axon calibers. We found that 75.15 ± 3.15% of axons had a caliber smaller than 1.1 µm, likely causing the net increase in myelin content observed in MRI data (**Fig. 8m,n**). In summary, these findings demonstrate that CRH/CRHR1 signaling is not only activated under injury conditions, but plays an important role in regulating OL generation during early postnatal stages and, furthermore, influences adult myelination patterns by enhancing myelin sheet thickness in small caliber axons.

## Discussion

Although, the process of oligodendrogenesis and myelination is one of the main functions of OPCs, the modulation of this process is not fully understood. Neuropeptides, like CRH, represent a large and diverse group of molecules commonly considered as neuromodulators, which have been implicated in a wide variety of processes in the CNS. So far, they have been investigated predominantly in the context of neuronal development and as modulators of synaptic transmission (Guillaumin and Burdakov, 2021; Russo, 2017; van den Pol, 2012). In our study, we identified and characterized a novel CRH/CRHR1 system in the OL lineage. We found that this system modulates OPC differentiation and subsequent myelination following injury, as well as under non-injury conditions. Initially, we discovered a subpopulation of CRH-expressing OPCs constituting 30% of the total population of injury-responsive OPCs. This contribution distinguishes them from other OPC subpopulations, such as the GPR17^+^ OPCs, which only contribute ≈14% to the whole OPC population and underscores their potential impact on the general injury response (Miralles et al., 2023). CRH expression was observed to commence within 12 hpi, a finding confirmed by a published single cell sequencing dataset (Li *et al*., 2022). The identification of CRH expression in OPCs not only represents the first comprehensive *in vivo* description of neuropeptide expression in OPCs but also one of the earliest reported responses of OPCs to acute injury. Although their population dynamics are similar to the general population of OPCs, CRH-expressing OPCs differ in their specific proliferative response as well as their differentiation tendency. Proliferation is markedly increased (100% proliferation rate) in this population compared to all OPCs responding to cortical injury (40%) (von Streitberg *et al*., 2021).

Additionally, their high maturation rate of 80% distinguishes them from the general population of cortical OPCs, which have been reported to possess a relatively low oligodendrogenic potential (Dimou *et al*., 2008; von Streitberg *et al*., 2021). These discrepancies were not caused by regional differences in the OPC population between CX and MB, as shown by the comparison to other MB OPCs, which clearly highlights that CRH-expressing OPCs exhibit a faster and higher differentiation rate. Additionally, their significant role in injury-related recovery is emphasized by their extensive contribution to the population of newly generated OLs (≈40%) and their remarkable stability after integration, as evidenced by the persistence of OLs derived from CRH-expressing OPCs up to four months following acute injury (Tripathi *et al*., 2017).

Given the absence of CRH-expressing OPCs in the brain under non-injury conditions, we first investigated their role in regenerative processes following injury. Our findings indicate that OPC-derived CRH delays the differentiation of another OPC subpopulation expressing CRHR1, thereby, facilitating long-term survival of OLs. This hypothesis is supported by the identification of CRHR1-expressing OPCs surrounding the injury site, which also generate myelinating OLs. While CRHR1 expression in OPCs was confirmed on different levels in this study, its expression has also been identified in different sequencing studies, further strengthening our finding (Brown et al., 2016; Falcao et al., 2019; Zeisel et al., 2018; Zeisel et al., 2015; Zhang et al., 2014). Following injury, we found that the dynamics of oligodendrogenesis in the CRHR1^+^ OPC population are slower compared to the whole OPC population. Furthermore, CRHR1 inactivation results in an increase in the differentiation velocity, leading to a long-lasting loss of OLs in an injury size-dependent manner. Therefore, CRHR1 seems to act in a similar way as GPR17, which has also been identified as a modulator of OL generation velocity (Chen et al., 2009). That premature differentiation can result in cellular loss before stable integration was already shown in other studies (Genoud et al., 2002). The necessity for the delay caused by CRH/CRHR1 interaction following injury may be related to the exposure to conditions of high oxidative stress present at early stages, which OLs, depending on their differentiation state, are particularly vulnerable to (Back et al., 1998). Evidence that an inflammatory environment can impact OPC differentiation was just recently provided in a study by *Meijer et al*, in which BCAS1 expression was epigenetically downregulated following treatment with Interferon γ (Meijer et al., 2022). Besides, other studies have already shown that timing of cell-cycle exit, subsequent OPC differentiation and initiation of maturation are tightly regulated by different factors (Barres et al., 1994; Chen *et al*., 2009; Palazuelos et al., 2014). It remains unclear why CRH-expressing OPCs seem to be less vulnerable to this environment and only prevent CRHR1-expressing OPCs from premature differentiation. One hypothesis is that there are intrinsic differences between the different OPC subpopulations that go beyond their expression of CRH and CRHR1.This is further indicated by the fact that CRH-expressing OPCs only occur under injury, but not under non-injury conditions, whereas CRHR1-expressing OPCs are always present. This raises the question of additional triggers of CRH-expression in OPCs but also for other sources of CRH, e.g. neurons, that might influence CRHR1-expressing OPCs and OLs.

When assessing the population dynamics of CRHR1-expressing OPCs under non-injury conditions, we found that they displayed a significantly elevated tendency for differentiation compared to other OPCs. Hence, when serving the same transient inhibitory purpose under physiological conditions as following injury, the inactivation of CRHR1 was suspected to further increase their oligodendrogenic potential. Indeed, we found that CRHR1 ablation lead to premature differentiation at early postnatal timepoints as well as to an increase in adult myelination as depicted by anti MBP staining, MRI-dependent myelin assessment and EM. This increase in the total amount of myelin was accompanied by a shift in the myelination pattern, with thicker myelin sheets in small-caliber axons and thinner sheets in large-caliber axons. This observation points in the direction that, although OLs seemingly myelinate small caliber axons prematurely and to a higher extent, their capacity to myelinate large caliber axons is impaired. The question why CRH either from OPC or non-OPC sources has such a remarkable influence on myelination remains elusive, but might be connected to its physiological role as a stress-responsive neuropeptide (Chang et al., 2022; Peng et al., 2017). Previous studies have shown that stress, particularly during the early postnatal phase, can disrupt oligodendrogenesis, leading to long-lasting effects on both myelin structure and animal behavior (Bonnefil et al., 2019; Bordner et al., 2011; Breton et al., 2021; Raabe et al., 2019; Teissier et al., 2020; Yang et al., 2016). Given that CRH is widely recognized as a stress peptide involved in stress-related pathologies, exploring its impact on the CRH/CRHR1 system in OPCs is a natural direction for future research (Nishiyama *et al*., 2021). The fact that neuropeptides, unlike fast-acting neurotransmitters, possess slow release kinetics and primarily act through GPCRs via volume transmission, makes them ideally suited to modulate long-lasting effects on myelination (Fields, 2004; van den Pol, 2012). Ultimately, this study unravels i) a previously unknown context-dependent secretory response of the neuropeptide CRH by OPCs, ii) its direct impact on the myelinogenic potential of a second, CRHR1-expressing, OPC subpopulation, and iii) that CRHR1-expressing OPCs, regardless of the CRH source, exhibit an increased oligodendrogenic tendency, which is further enhanced by CRHR1 inactivation, leading to altered myelin structure. Together, these findings suggest that CRH – and potentially other neuropeptides – can significantly influence oligodendrogenesis and myelination under physiological and pathological conditions.

## Online Methods

### Animals

All animal experiments and protocols were legally approved by the Ethics Committee for the Care and Use of Laboratory Animals of the Government of Upper Bavaria, Germany. 2-6 month old mice were group housed under standard lab conditions (22 ± 1°C, 55 ± 5% humidity) with ad libitum access to food and water on a 12:12 h light:dark schedule with weekly cage changes. Regular genotyping was performed by polymerase chain reaction (PCR) analysis of tail DNA. Global and conditional knockouts and global and conditional overexpressing mice were assigned to experimental groups based on their genotype. Age matched littermates were used as controls in all experiments. Mice with incorrect injury or injection were excluded from the experiment. The following mouse lines were used:

*CRH-Cre* (*Crh^tm1(cre)Zjh^*, Jackson Laboratory stock no. 012704)(Taniguchi et al., 2011), *CRH-FlpO* (*Crh^tm1.1(flpo)Bsab^*, Jackson Laboratory stock no. 031559)(Salimando et al., 2020), *CRHR1-Cre* (*Crhr1^tm4.1(cre)Jde^*)(Dedic et al., 2018), *CRH-Venus* (*Crh^tm1.1Ksak^*)(Kono *et al*., 2017), *Tau-LSL-FlpO(Pivetta et al., 2014)*, *NG2-CreERT2* (*Tg(Cspg4-cre/Esr1*)BAkik*, Jackson Laboratory stock no. 008538)(Zhu et al., 2011),, *Ai9* (*Gt(ROSA)26Sor^tm9(CAG-tdTomato)Hze^*, Jackson Laboratory stock no. 007909)(Madisen et al., 2010), *Sun1-GFP* (*Gt(ROSA)26Sor^tm5(CAG-Sun1/sfGFP)Nat^*, Jackson Laboratory stock no. 021039)(Mo et al., 2015), *Ai65* (*Gt(ROSA)26Sor^tm65.1(CAG-tdTomato)Hze^*, Jackson Laboratory stock no. 021875)(Madisen *et al*., 2010), *Ai65F* (*Gt(ROSA)^26Sortm65.2(CAG-tdTomato)Hze/J^*, Jackson Laboratory stock no. 032864*), CRH^loxP^* (*Crh^tm1.1Jde^*)(Dedic *et al*., 2018), *CRHR1^loxP^* (*Crhr1^tm2.2Jde^*)(Kuhne et al., 2012), *CRHR1^ΔEGFP^* (*Crhr1^tm1Jde^*)(Refojo *et al*., 2011).

The following double and triple transgenic lines were generated in this study by cross-breeding of single transgenic lines (detailed description included in Table S1): *CRH-Cre::Ai9*, *NG2-CreERT2::Ai9*, *CRH-FlpO::NG2-CreERT2::Ai65*, *CRH-FlpO::CRHR1-Cre::Ai65*, *CRH^NG2-cKO^ (CRH^loxP^::NG2-CreERT2:: Ai9*), *CRHR1-Cre:Tau-LSL-FlpO::Ai9*, *CRHR1-Cre::Sun1-GFP*, *CRH-FlpO::Ai65F::CRHR1-Cre::Sun1-GFP*.

### Stereotaxic surgeries

For all experiments requiring stereotaxic surgeries, mice received analgesic treatment prior, during and after surgery. Animals were anesthetized using isoflurane (CP-Pharma) and placed in a stereotaxic frame. Acute injury, fluorescent bead, CRH, saline and virus injections, cranial window or hippocampal cannula implantation was performed as described in the sections below.

### Acute injury for CRH^+^ OLC quantification and modulation experiments

After opening of the skin and removal of a bone flap using a dental drill, the syringe was slowly inserted and removed from the tissue. The following coordinates were used: PFC: AP 2.2, ML 1.0, DV -3.2; Striatum: AP 1.2, ML 1.5, DV -3.3; MB: AP -3, ML 1.0, DV -4. Injuries were inflicted using a 24-gauge Hamilton syringe. Injuries for label retaining experiments were performed using a 33-gauge Hamilton syringe to reduce background for BrdU staining.

### Fluorescent bead-, CRH-, saline-and CMV-GFP-virus injection

For injections, head skin was opened. A bone hole was drilled using a dental drill. The syringe was inserted. Then, Fluorencent beads (Green Retrobeads™ IX, Lumafluor, 500 nl), or CMV-eGFP-virus (AAV1/2-CMV-DIO-eGFP, Vector Biolabs, 500 nl) were injected at a speed of 100 nl/ min with a 33 gauge syringe (Hamilton). After injection was finished, the syringe was pulled out 1 mm and, after a waiting time of 2 minutes, slowly and finally removed from the tissue. The following coordinates were used: AP -3, ML 1.0, DV -3.7.

### Cranial window implantation with injury infliction

After stereotaxic fixation, a round piece of skin, ∼8 mm in diameter, was removed. Under repeated application of cold saline to prevent overheating, a round piece of cranial bone was drilled out (approx. 5 mm diameter). A double-edged knife was moved in (DV: -0.8 mm) and out of the brain parenchyma three times. A drop of sterile saline was applied to the open brain tissue and a 5 mm cover slip was placed over the opening and subsequently fixed using quick adhesive cement (Parkell C&B Metabond clear powder L, Quick Base B, Universal Catalyst C). The custom-made head plate was positioned on top of the head and fixed using Kallocryl (Speiko, liquid component 1609 and powder 1615).

### Implantation of subcortical imaging cannula

Preparation and implantation of the subcortical imaging cannula were performed as described before(Ulivi et al., 2019). The cannula consisted of a cylindrical metal tube, 1.6 mm height and 3.5 mm diameter, sealed at the bottom with a glass coverslip. After stereotaxic fixation of the animals, the head skin was removed. Under repeated application of cold saline to prevent overheating, a round piece of cranial bone (approx. 3.5 mm diameter) was gently carved using a trephine. After removal of the bone flap, the cortex was slowly aspirated using a vacuum pump until the corpus callosum, identified as white matter horizontal fibers, was exposed. The imaging cannula was gently inserted into the cavity and immediately fixed using a quick adhesive cement (Parkell C&B Metabond, clear powder L, Quick base B, Universal Catalyst C). Finally, a custom-made head plate was fixed as described above.

### In-vivo 2-photon imaging

2-photon imaging experiments were done using an Ultima IV microscope from Bruker, equipped with an InSight DS+Dual laser system. We used a 1040 nm to excite tdTomato, while we tuned the laser to 920 nm wavelength to image the dura. During imaging, the average power at the tissue did not exceed 40 mW. Animals were anesthetized using isoflurane and fixed under the microscope using a custom-made head plate holder. We kept the animals’ temperature constant by means of a heating pad set to 37 °C. Z-stacks were acquired with 2-5 µm step size using an Olympus XLPlan N 25×/1.00 SVMP objective at magnification zoom 1x and 2.38x. We acquired images averaged over 4-16 repetitions per each imaging plane. For time-lapse imaging, images were acquired every 60 s for 20-30 min.

### Magnetic resonance imaging (MRI)

MRI experiments were run on a BioSpec 94/20 animal MRI system (Bruker BioSpin GmbH) equipped with a 9.4 T horizontal magnet of 20 cm bore diameter and a BGA12S HP gradient system capable of a maximum gradient strength of 420 mT/m with a 140 µs rise time. Each mouse in the cohort was sedated in a preparation box using 2.5 vol % isoflurane. The animal was then fixed in prone position on an MR compatible animal bed using a stereotactic device, while anesthesia was delivered via a breathing mask. Mice were kept anesthetized with an isoflurane/air mixture (1.5–1.8 vol %, with an air flow of 1.2–1.4 L/min). Respiration and body temperature were constantly monitored using a pressure sensor placed below the mouse’s chest and a rectal thermometer, respectively. Body temperature was kept between 36.5 °C and 37.5 °C using a heating pad. A linear volume resonator coil for excitation of the ^1^H nuclear spins in combination with a 2x2 elements surface array radio frequency coil for reception of their signal were used. After positioning of a mouse in the magnet isocenter, a field map-based shimming was performed to optimize B_0_ field homogeneity over the entire mouse brain. Structural T2 –weighted (T2w) images were acquired using a 2D multi-slice RARE sequence (RARE factor= 8, TE_eff_/TR = 33/2500 ms.

### Magnetization transfer (MT) to assess Molecular Proton Fraction (MPF)

Magnetization transfer (MT) methods in MRI can be used to estimate the Macromolecular Proton Fraction (MPF) which has been shown to be a sensitive biomarker of myelin density (Hertanu *et al*., 2023; Khodanovich *et al*., 2017; Soustelle *et al*., 2021; Yarnykh, 2012; Yarnykh, 2016). The protocol was derived from recommendations given in Soustelle et al. 2021 and Hertanu et al. 2023(Hertanu *et al*., 2023; Soustelle *et al*., 2021). First, four averages of an MT prepared spoiled gradient recall experiment (MT-SPGR) were acquired, using a preparation Gaussian saturation rf-pulse of 10.25 ms at an offset frequency of 6 kHz and a flip angle of 600°, resulting in a B_1peak_ of 9.1µT (Soustelle *et al*., 2021). This pulse was followed by an SPGR with an echo time TE of 2.2 ms and a recovery time TR of 30 ms. The matrix size for the experiment was 128 × 128 × 64, with a field of view of 15 × 15 × 25 mm^3^, resulting in a resolution of 0.1 × 0.1 × 0.39 mm^3^. The duration of the experiment was 14 minutes and 20 seconds. Second, variable flip angle (VFA-) SPGR sequences were performed for T1 (R1) quantification. Three such sequences were recorded using flip angles of respectively 6°, 10° and 25°. TE, TR, matrix size, field of view, resolution and number of averages for each of the three VFA-SPGRs were the same as for the MT-SPGR, resulting in the same duration for each of them. Third and last, a B_1_^+^ mapping experiment was recorded with the actual flip angle imaging (AFI) method (Yarnykh, 2007).TE was 2.388 ms, TR 15 ms and the flip angle was 60°. The matrix size was 44 × 44 × 40, the field of view 15 × 15 × 25 mm^3^ resulting in a resolution of 0.341 × 0.341 × 0.625 mm^3^. With a number of averages of four, the duration for the experiment was 6 min 37 s.

Based on these (MT-, T1 and B_1_^+^) experiments plus the B_0_ map obtained from the field map recorded in order to perform the map-based shimming, our data could be prepared to use the software freely available at https://github.com/lsoustelle/qMT in order to derive the MPF for each individual animal (Hertanu *et al*., 2023; Hertanu et al., 2022; Soustelle et al., 2020). From these individual maps, an MPF template could be derived using routines in the Advanced Normalization Tools (ANTs) software (http://stnava.github.io/ANTs/). First a few representatives, manually aligned, individual MPF maps were co-registered and merged together into a starting template. Then the routine antsMultivariateTemplateConstruction2.sh in the ANTs software was used to build a template based on all the MPF maps corresponding to each animal in the cohort. Each individual map could then be, in turn, normalized to the newly created template.

### Experimental setup modulation of CRH/CRHR1 system in glial cells

For experiments with inducible Cre recombinase lines, recombination was induced by i.p. injection of TAM two times within the 7 days before injury infliction. For modulation experiments, animals were injured or injected as described above. Mice were sacrificed at 3 or 7 dpi and analyzed for number of OLs (CC1^+^/ Olig2^+^), OPCs (CC1^-^/ Olig2^+^), all OLCs (Olig2^+^) cells and the percentage of OLs of all OLCs (CC1^+^ of all Olig2^+^ cells).

### Label retaining experiment

BrdU 1 mg/ kg was added to the drinking water (1 % w/w sucrose). In case of injury experiments, treatment started at the day of injury infliction and was sustained for a maximum of 7 days (7 dpi and 6 wpi mice) or until sacrifice (3 dpi). For naive labeling of CRHR1 KO mice, treatment was applied for two consecutive weeks followed by a retaining time of three weeks.

### Cryosectioning

Animals were sacrificed using isoflurane and subsequently perfused with ice cold 1xPBS and 4 % PFA. After recovery of the brain, it was post fixed in 4 % PFA on ice for 6 h, transferred to 30 % sucrose in 1xPBS. and incubated at 4 °C for 48 h. Brains were frozen on dry ice and cut either coronally or horizontally in 40 µm sections using a cryostat (Leica). Sections were collected in cryoprotection solution (25 % Ethylene glycol, 25 % glycerol, 50% dH2O in 1xPBS) and stored at -20 °C until further use. For DISH mice were killed by cervical dislocation. After fast recovery the brain was frozen on dry ice and stored at -80 °C until further use.Coronal sections (20 µm) were generated using a cryostat (Leica). After thaw-mounting onto SuperFrost slides, sections were dried and kept at -80 °C.

### Immuno fluorescence staining

Immuno fluorescence staining was conducted using different protocols depending on the combination of antigens of interest. For NG2/ Olig2 and NG2/Ki67 staining, a two-day protocol was performed. In brief, slices were washed 3x in 1x PBS, followed by blocking in 2 % normal goat serum in 0.05 % Triton-X100 and 1x PBS. Sections were incubated in primary anti-NG2 antibody at 4 °C under shaking overnight. After washing, sections were incubated in secondary antibodies for 2 h at room temperature (RT). After washing, antigen retrieval (AR) was performed in citrate buffer (75 °C, 1 h). After washing, primary anti-Olig2 or anti-Ki67 antibody was incubated as described before. After washing, secondary antibody was applied as described before and sections were washed and mounted using Fluoromount-G^TM^ mounting medium (+/- DAPI, Invitrogen, 15586276).

For CC1/ Olig2, Ki67/ Olig2 and Ki67/ Olig2/ GFP staining, washing was performed followed by AR as described before. After washing, sections were blocked. Primary and secondary antibody incubation were also conducted as described before.

For anti-GFP, anti-GFAP, anti-Vimentin, anti-S100β, anti-IBA1, anti-NeuN, anti-MBP, anti-CNPase and anti-CD31 staining sections were washed and blocked as described above. Primary and secondary antibody treatment were performed as mentioned above.

For CRH, staining sections were washed and blocked as described before. Primary antibody was added and sections were incubated for 5 days under shaking at 4 °C. Secondary antibody treatment was performed as described above.

For BrdU staining, two different protocols were used. Sections were washed and AR was achieved either with citrate buffer (2.94 g/l, pH 6, 1 h, 75 °C) for injury sites or with HCl (2 M, 10 min, 37 °C) followed by borate buffer (10 min, RT) for naive brains. After washing sections were blocked and primary and secondary antibody staining was performed as described above.

For GPR17 staining sections were quenched with H_2_O_2_ (1 % in dH_2_O) for 15-30 min. Sections were blocked and incubated with anti-GPR17 Ab as described above. Secondary antibody staining was performed using a biotinylated goat-anti-rabbit (rb) Ab for 2 h at RT. After washing horse-radish-peroxidase (HRP) was applied for 30 min at RT. Samples were incubated in Tyramine solution (stock 1:20 in 1x amplification dilution) for 3-10 min. After washing samples were mounted and covered as described above.

The following primary antibodies and dilutions were used: rb NG2 (1:200, Merck, AB5320), rb Olig2 (1:200, Merck, AB9610), mouse (ms) Olig2 (1:200, Merck, MABN50), ms APC (CC1) (1:200, Merck, OP80), rb GPR17 (1:500, Sigma, SAB4501250), ms CNPase (1:1000, Abcam, ab6319), rb MBP (1:500, obtained from Klaus-Armin Nave, MPI exp. Medicine, Goettingen)(Meschkat et al., 2022), rat MBP (1:500, Merck, MAB386), ms MBP (1:200, Biolegend, 836504), rb Ki67 (1:500 - 1:1000, Abcam, Ab15580), rat BrdU (1:500 – 1:1000, Abcam, 6326), rb Iba1 (1:500 – 1:1000, Synaptic Systems, 234013), ms NeuN (1:500, Abcam, Ab13970), rb GFAP (1:500 – 1:3000, Abcam, ab7260),,, rb CRH (1:20000, obtained from Paul E. Sawchenko, Salk Institute, CA.), ck GFP (1:500 – 1:1000, Aves, GFP-1020), rb RFP (1:500, Rockland, 600-401-379),.

The following secondary antibodies were used: goat anti rb Alexa 488, Alexa 568, Alexa 594 or Alexa 647, goat anti ms Alexa 488, Alexa 568, Alexa 594 or Alexa 647, goat anti rat Alexa 488 or Alexa 568 and goat anti ck Alexa 488.

For expansion staining following expansion microscopy the following secondary antibodies were used: anti ms STAR RED, Abberior Cat# STRED-1001-500UG, anti rb STAR ORANGE, Abberior Cat# STORANGE-1002-500UG.

### Expansion microscopy

Brain sections of *CRHR1-Cre::LSL-FlpO::Ai9* mice were retrieved from cryoprotectant solution and washed three times with 1x PBS. To expand the samples, an adjusted TREx protocol was applied, as previously described^(Damstra^ ^et^ ^al., 2022; Delling^ *^et^ ^al.^*, ^2024)^. Briefly, brain slices were mounted onto super-frost microscope slides and treated with 10 µg/mL acryloyl X-SE in 1x PBS overnight at RT. The gelation solution contained 1.1 M sodium acrylate, 2.0 M acrylamide, 50 ppm N,N′-methylenebisacrylamide and 1x PBS. The polymerization was initiated by the addition of TEMED (1.5 ppt) and APS (1.5 ppt). The process was slowed by 4-Hydroxy-TEMPO (15 ppm) and working on ice to allow for an extended incubation time of 30 min in activated gelation solution, before mounting the slide containing the brain sections into a custom-build gelation chamber plate. The samples were incubated for 2 h at 40 °C to allow for full polymerization. Then, slices were recovered into digestion buffer (50 mM Tris-BASE, 200 mM NaCl, 200 mM SDS in ddH_2_O, pH 9.0) and incubated for 4 h at 80 °C in a Thermo-Block. After denaturation, the samples were rinsed once in ddH_2_O and washed several times in 1x PBS to remove residual SDS. At this stage, the region of interest was cut from the gel with a razor blade. For immune fluorescent staining, the samples were incubated for 3 h in blocking solution (0.3% Triton-X-100, 3% BSA in 1x PBS), followed by 72 h of incubation with primary antibodies (ms MBP, rb RFP) diluted in blocking solution at 4 °C under shaking. Subsequently, secondary antibody incubation (ms STAR RED, rb STAR ORANGE) was performed overnight at 4 °C under shaking. The stained gels were then incubated with BODIPY-FL NHS (20μM, ThermoFisher) in 1x PBS for 1h at RT on a shaker. Afterwards, the samples were washed five times in ddH_2_O and stored at 4 °C overnight to complete the expansion of the gel. For imaging, the gels were placed onto the PLL-coated surface of a #1.5H one-chambered cover glass (Cellvis), sample-side facing down. The samples were then immobilized by embedding them into two-component silicone (eco-sil, picodent), preventing drift and dehydration during imaging.

### Image acquisition

Images for quantification of cell numbers and intensity measurements were acquired using an Olympus VS120 Slide Scanner. For overview and close up acquisition, the 4x or 20x objective was used, respectively. Exposure times were chosen to yield the best signal to noise ratio and minimal photodamage. Images were extracted and saved as .tif files.

Qualitative images of cells of interest were taken with a Laser scanning confocal microscope (Carl Zeiss). 20x water-, 40x and 63x oil-immersion objectives were used. Laser settings were adjusted to yield the best signal to noise ratio. Images were acquired with 1024x1024 pixel size and a scan speed between 3 and 7.

### Quantification of CRH-expressing OPCs in *CRH-Cre::Ai9* mice

A 28 µm z-stack with a step size of 3 µm was acquired around the injury site. The counting matrix was generated with a custom Fiji macro. Cells in the different areas between +/-300 µm were counted using Fiji’s manual counting tool. To generate comparable counts, consistent contrast settings were used for each quantification.

Quantification of CRHR1-expressing OLCs at 1.5, 3 and 5 months in *CRHR1-Cre::LSL-FlpO::Ai9* mice. After CC1/ Olig2 or NG2/ Olig2 staining was performed, z-stacks (28 µm depth, 3 µm step size, 1 mm^2^ ROI) were acquired in CX, CC, AC, OT and MB. CC1 and NG2/ tdTomato co-expressing cells were quantified in the whole ROI. The number of all Olig2^+^ cells was counted in a consistent subregion (64000 µm^2^) and upscaled to cells/ mm^2^.

### Quantification of cells in CRH/ CRHR1 system modulation experiments

Horizontal sections were imaged as described above. After setting the wound center, the circular counting matrix with a radius of 300 µm and medio-lateral resolution of 50 µm was generated using a custom macro in Fiji (**Method S1**). In the increased injury size experiment coronal sections were analyzed to increase the visible section of the injury. These injuries were analyzed using the rectangular counting matrix with a medio-lateral resolution of 50 µm which was semi-automatically created using a custom Fiji macro. Because of variations in the visible part of the wound and, hence, the height of the analyzed ROI the numbers are presented as cells/ mm^2^. Cells were counted with the manual Fiji counting tool. Consistent contrast settings were used for each quantification.

### MRI analysis

MPF images were analyzed in SPM12 (www.fil.ion.ucl.ac.uk/spm/software/spm12/) using a full factorial model with factors sex (male and female) and genotype (+/+ and lox/lox). For each of the four groups, n=8 animals were included in the final analysis. The whole brain analysis, excluding the CSF space using an implicit intensity mask, was run without global normalization, as the MPF is an absolute measure. F-contrasts of the main effects of sex or genotype were calculated, along with the interaction sex × genotype. To detect statistical differences, a post-hoc t-test showing the negative effect of genotype with a p_FDR,cluster_<0.05, at a collection threshold of p < 0.01 was conducted.

### Electron microscopy

Samples were prepared according to Weil and colleagues(Weil et al., 2019). In brief, animals were sacrificed using isoflurane and perfused with 1x PBS (pH 7.4) and subsequently with fixative solution (4% PFA, 2.5% glutaraldehyde in phosphate buffer with 0.5 % NaCl, pH 7.4). Brains were dissected and postfixed in same fixative overnight at 4°C. For targeting the region of interest, sagittal vibratome slices were prepared with a Leica VT1200S (Leica, Wetzlar, Germany) and pieces of medial and lateral rostral corpus callosum were extracted from the vibratome slices using a biopsy punch. These pieces were postfixed with 2 % OsO4 (Science Services, München, Germany) in 0.1 M phosphate buffer pH 7.3 and embedded in EPON resin (Serva, Heidelberg, Germany) after dehydration with acetone. Ultrathin sections were prepared using a Leica UC7 ultramicrotome (Leica, Wetzlar, Germany) and a 35° diamond knife (Diatome, Biel, Switzerland) and stained with UranylLess™ (Science Services, Munich, Germany). EM pictures were taken with a Zeiss EM912 electron microscope (Carl Zeiss Microscopy GmbH, Oberkochen, Germany) using an on-axis 2k CCD camera (TRS, Moorenweis, Germany).

### G ratio measurement and analysis

Image analysis was performed with ImageJ (Fiji, Version 2.0.0-rc-69/1.52). For g ratio analysis, 12 random overview EM pictures (at 5000x magnification) corpus callosum sections were taken and 300-350 axons per animal were analyzed. For g ratio analysis (axon diameter divided by the axon diameter including the myelin sheath) diameters were determined from circular areas equivalent to the measured areas. The analysis was carried out blinded with regard to the genotype.

### Statistical analysis

For statistical analysis, GraphPad Prism 8 was used. One-way ANOVA followed by Bonferroni’s and two-way ANOVA followed by Sidak’s post hoc test were performed. Values are reported as means ± standard error of mean (s.e.m.). Statistical significance was defined as P < 0.05. No statistical methods were used to predetermine sample sizes, but it was based on those previously reported in other publications(Dimou *et al*., 2008; Osso *et al*., 2021). During analysis, experimenters were blinded to experimental conditions.

## Data availability

Additional data, supporting the findings of this study are available from the corresponding author upon reasonable request.

## Code availability

All macros used in this study are available from the corresponding authors upon reasonable request.

## Acknowledgments

The authors thank Sabrina Bauer, Stefanie Unkmeir and Andrea Parl for their constant technical assistance and Elfi Fesl for proofreading and editing of the manuscript. We thank Sandra Goebbels for her scientific input and for sharing different antibodies with us. Additionally, we thank Mathias Schmidt for providing AAV1/2-CMV-DIO-eGFP virus. Two icons (Fig. 3a, Fig. 8h) were created with BioRender (BioRender.com). The project was supported by the Max Planck Society (JMD). CR received funding from the International Max Planck Research School for Translational Psychiatry (IMPRS-TP) and the Gunter Sachs Donation.

## Author Contributions

C.R. designed, led and performed all experiments, analyzed and interpreted data and wrote the manuscript. T.S. performed MRI measurements and analysis, provided essential scientific input and subedited the manuscript. B.B. setup the pipeline for MPF measurements, aided scientific input and subedited the manuscript. T.R. acquired EM pictures, provided essential scientific input on image analysis and subedited the manuscript. JP.D. performed tissue expansion, subsequent staining and subedited the manuscript. T.M.I. performed stainings and analyses, provided scientific input and subedited the manuscript. J.P. analyzed data and subedited the manuscript. A. U. helped with cannula implantations for *in vivo* imaging, gave scientific expertise, helped with image acquisition at the 2-photon setup and subedited the manuscript. S. C. aided methodological and scientific expertise. I.M.H. aided methodological expertise in cranial window implantation, provided scientific input and subedited the manuscript. K.S. and K.I. created and provided the CRH-Venus mouse line and reviewed the manuscript. D.N. provided material and funding for tissue expansion and subedited the manuscript. A.A. provided scientific input concerning 2-photon imaging, reviewed and subedited the manuscript. M.C. set up the imaging protocol and performed MRI, analyzed the data, provided valuable input on its interpretation and subedited the manuscript. K.A.N. provided resources for EM imaging, provided scientific advice and subedited the manuscript. W.M. was critically involved in the planning of the EM experiment, provided advice concerning its analysis and interpretation and subedited the manuscript. L.D. provided constant scientific input, helped with planning label retaining experiments, reviewed and subedited the manuscript. J. M. D. provided essential scientific advice and support, supervised all experiments and analysis, reviewed and edited the manuscript.

## Conflict of Interest

The authors declare no competing interests.

## Notes

### Competing Interest Statement

The authors have declared no competing interest.

