## Supplementary info for "Neuropeptide CRH prevents premature differentiation of OPCs following CNS injury and in early postnatal development"

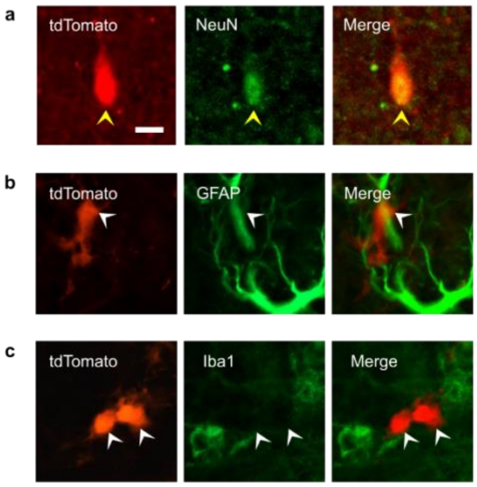


**Figure S1: Identification of CRH-expressing cells at injury. a-c**, Immuno stainings for NeuN, GFAP and Iba1 at the injury site in *CRH-Cre::Ai9* mice, showing only co-localization of tdTomato with NeuN. For all images, White arrowheads indicate cells or structures. Yellow arrowheads indicate co-localization of two markers.

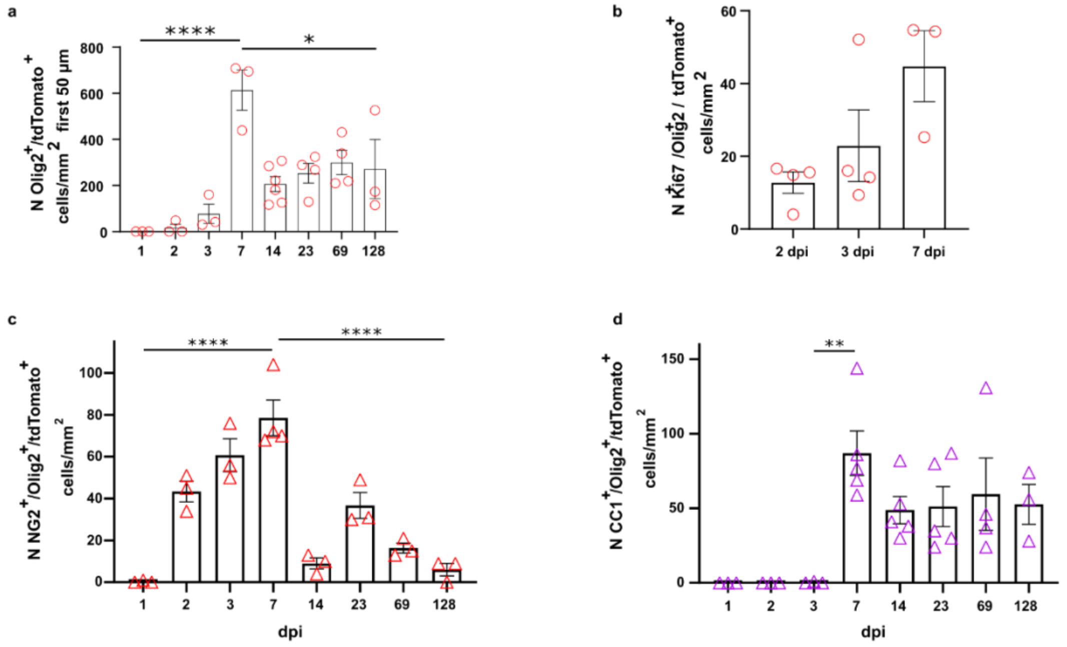


**Figure S2: Population dynamics and proliferation of CRH$^+$ OPCs following acute injury. a**, Quantification of Olig2$^+$/tdTomato$^+$ cells at ± 50 µm around the injury site. Changes were observed over time (One-way ANOVA: $F_{(7, 21)} = 10.27$, $p < 0.0001$) with an increase between 1 and 7 dpi ($p < 0.0001$, 95% Confidence interval (C.I.) = -909.2, -318.7) followed by a decrease between 7 and 128 dpi ($p = 0.0158$, 95% C.I. = 47.3, 637.8). **b**, Quantification of Ki67$^+$/Olig2$^+$/tdTomato$^+$ cells/mm$^2$ at ± 300 µm around the injury site showing non-significant increase in the number of CRH-expressing Ki67$^+$ cells. $n_{TP}$ = 3 - 4 mice. **c**, Quantification of NG2$^+$/tdTomato$^+$ cells/mm$^2$ at ± 300 µm around the injury site. Significant changes in cell numbers over time were observed (One-way ANOVA: $F_{(7, 17)} = 26.14$, $p < 0.0001$) with a significant increase between 1 and 7 dpi ($p < 0.001$, 95% C.I. = -106.9, -49.48) followed by a decrease between 7 and 128 dpi ($p < 0.0001$, 95% C.I. = 43.81, 101.2). **d**, Quantification of CC1$^+$/tdTomato$^+$ cells/mm$^2$ at ± 300 µm around the injury site. Significant changes in cell numbers over time were observed (One-way ANOVA: $F_{(7,23)} = 5.186$, $p = 0.0012$ with a significant increase between 3 and 7 dpi ($p = 0.0081$, 95% C.I. = -158.4, -14.90) followed by a non-significant decrease. Between 14 and 128 dpi cell numbers were constant. $n_{TP}$ = 3 - 5 mice.

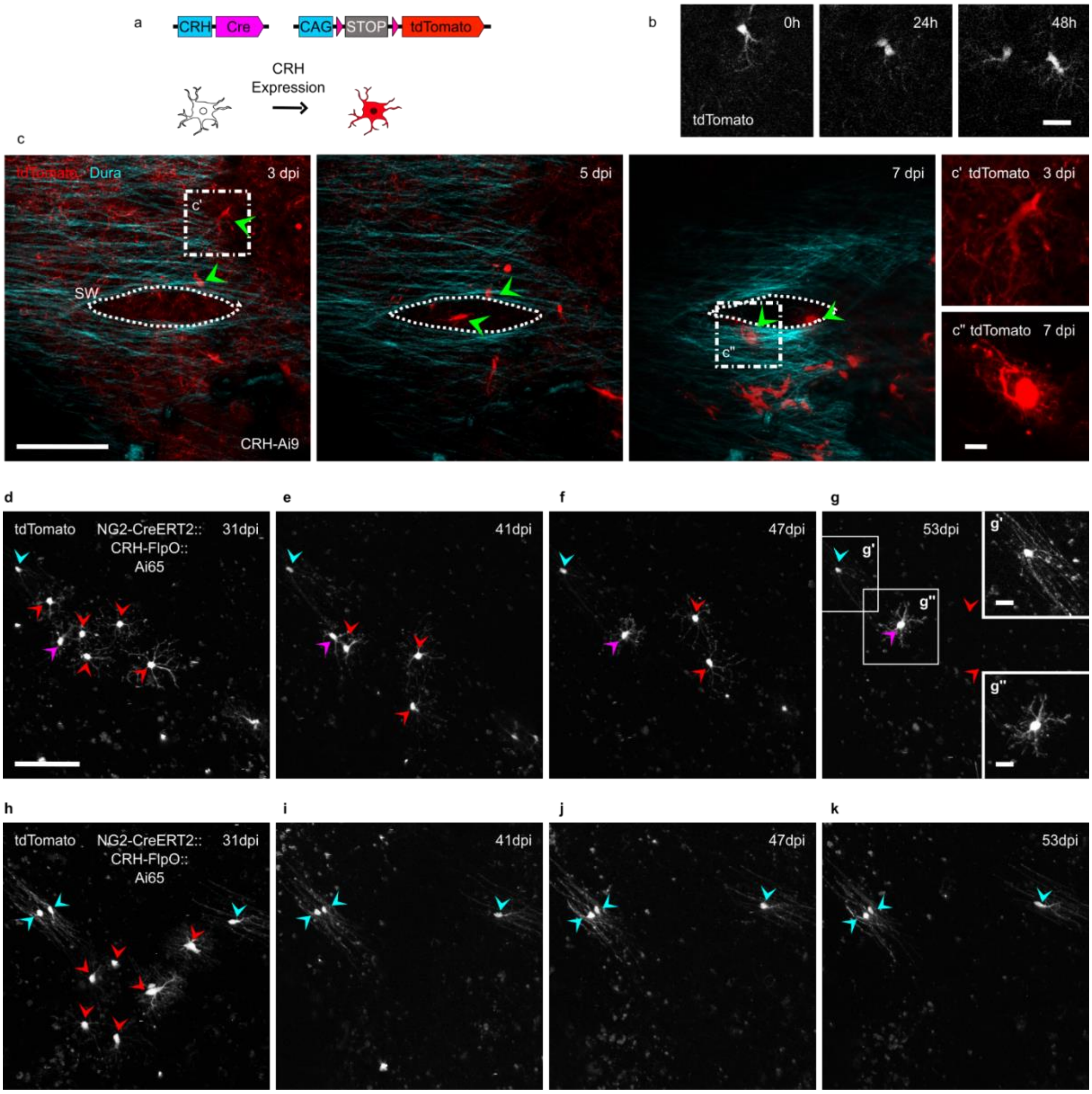


**Figure S3: CRH-expressing OLCs show disappearance without stable integration. a,** Graphical illustration of *CRH-Cre::Ai9* mouse model. **b,** 2-photon image of a proliferating tdTomato$^{+}$ OPC in WM over the course of 48 h. Scale bar, 20 µm. **c**, Representative 2-photon images of a cortical injury (white lining: injury site in dura) between 3 and 7 dpi. Arrowheads (green) indicate cells moving towards wound center. Scale bar, 100 µm. **c'** and **c''**, morphological change of single cell between 3 and 7 dpi. Scale bar, 10 µm. **d-k**, 2-photon *in vivo* images of CRH-expressing OPCs after hippocampal cannula implantation at 31 (**d, h**), 41 (**e, i**), 47 (**f, j**) and 53 dpi (**g, k**) showing disappearance of immature premyelinating OLs with highly motile processes and growth cone like structures and long-lasting stability of mature OLs. For all images: red arrowheads indicate disappearing cells. Purple arrowhead indicates persisting cells. Cyan arrowheads indicate stable, myelinating OLs (as judged by morphology). Scale bars, 100 µm (overview), 20 µm (close up **g' and g''**).

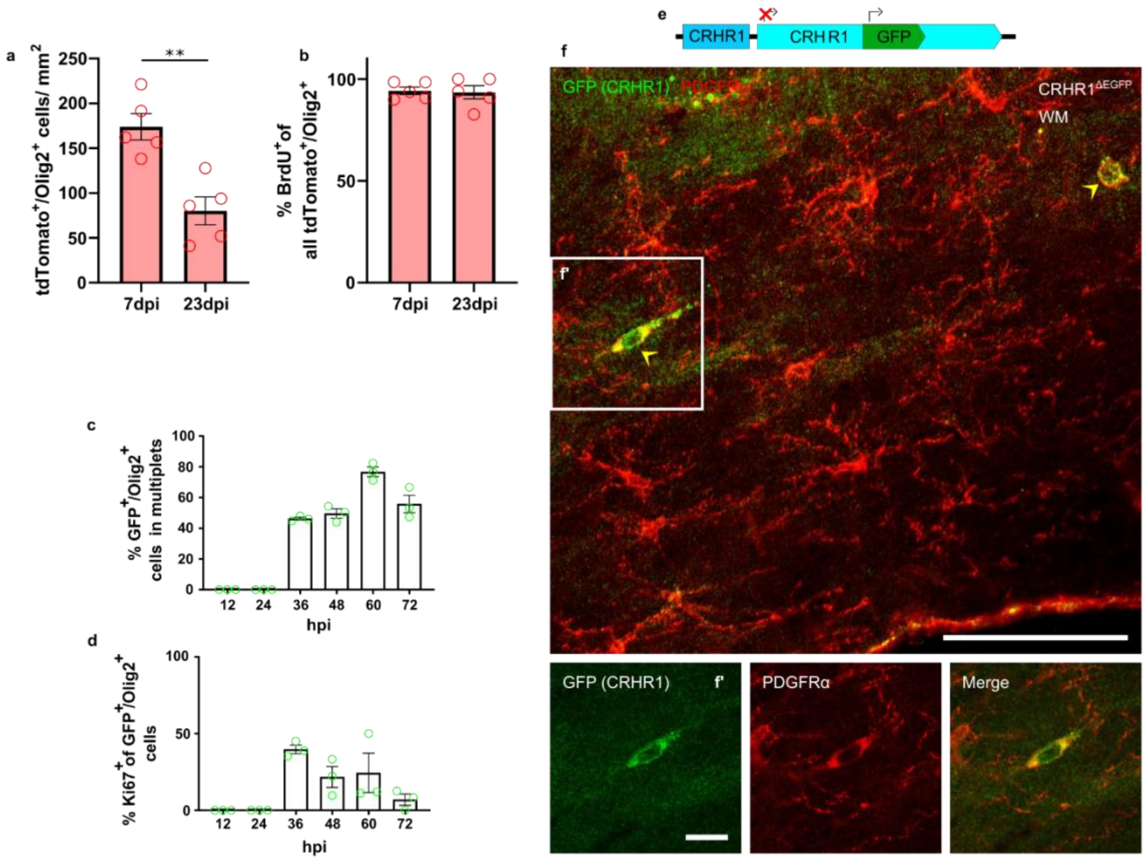


**Figure S4: Characterization and quantification of CRHR1-expressing OLCs**.

**a**, Number of tdTomato$^+$/Olig2$^+$ cells/mm$^2$ in *CRH-Cre::Ai9* mice showing a significant decrease from 7 to 23 dpi (Welch's two-tailed t-test: $t_{(8)} = 4.399$, $p = 0.0023$, $n_{TP} = 5$ mice). **b**, Percentage BrdU$^+$ of all tdTomato$^+$/Olig2$^+$ cells in *CRH-Cre::Ai9* mice at 7 (94.225 ± 1.83) and 23 dpi (93.49 ± 3.24). ***c***, Percentage of NG2$^+$/GFP$^+$ cells occurring in a multiplet of cells. $n_{TP} = 3$ mice. **d**, Quantification of Ki67$^+$ cells of all GFP$^+$/Olig2$^+$ cells. $n_{TP} = 3$ mice. **e**, Graphical illustration of CRHR1$^{\Delta EGFP}$ mouse model. **f**, GFP and PDGFRα co-staining in WM of CRHR1$^{\Delta EGFP}$ mice showing CRHR1$^+$/PDGFRα$^+$ cells. Scale bar, 100 µm (overview). 20 µm (close up **f'**). Yellow arrowheads indicate co-localization of two markers.

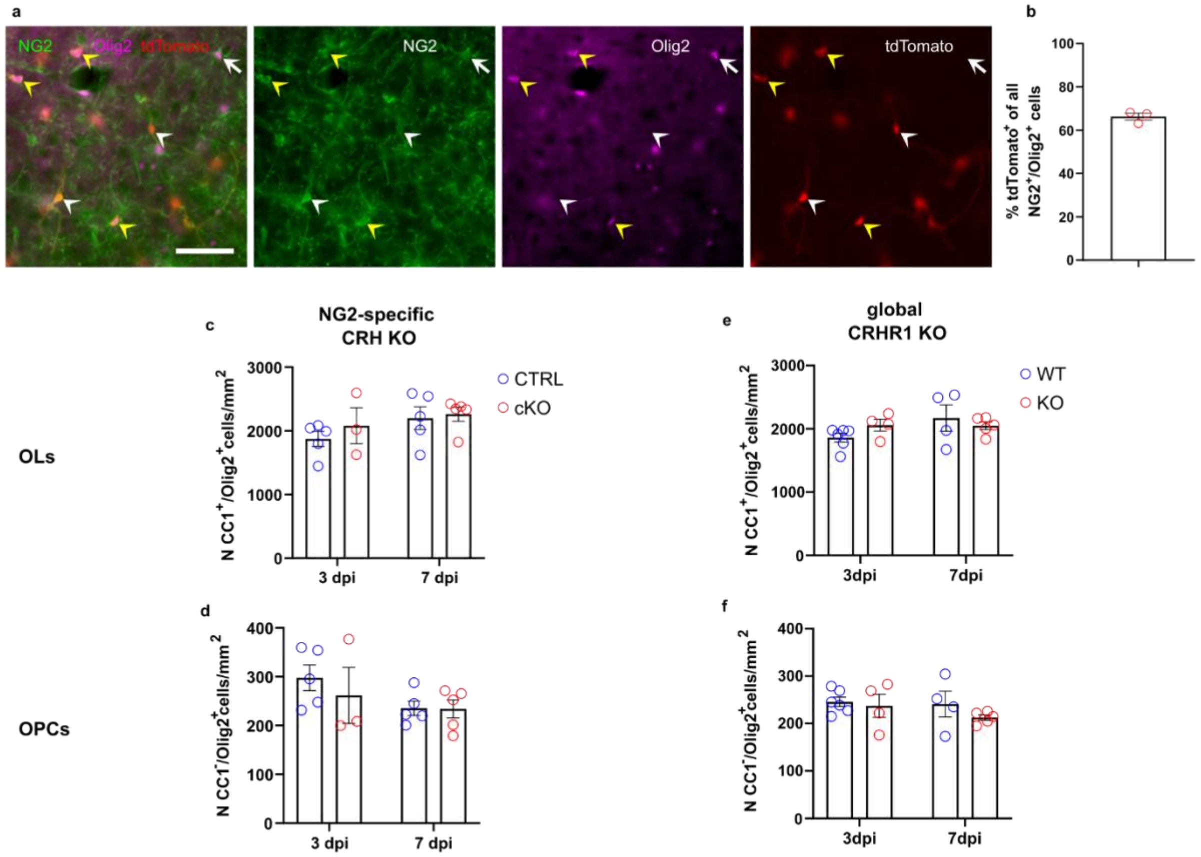


**Figure S5: Characterization of injury site in GOF experiments**. **a**, Representative image of tdTomato expression in NG2$^+$/Olig2$^+$ cells following TAM induction in NG2-specific CRH KO (CRH$^{NG2\text{-}cKO}$) animals showing effective Cre induction and consequent CRH KO. Scale bar, 50 µm. Arrowheads indicate recombined OPCs (tdTomato$^+$/NG2$^+$/Olig2$^+$) (yellow) and recombined pericytes (tdTomato$^+$/NG2$^+$/Olig2$^-$) (white), arrows indicate not-recombined OPCs (tdTomato$^-$/NG2$^+$/Olig2$^+$). **b,** Quantification tdTomato$^+$ cells of all NG2$^+$/Olig2$^+$ cells showing a recombination efficiency of 66.3 ± 1.6 %. **c-e**, Quantification of CC1$^+$/Olig2$^+$ (**c,e**), CC1$^-$/Olig2$^+$ (**d,f**) in non-injured region of NG2-specific CRH KO (**c,d**) and global CRHR1 KO (**e,f**) mice. Numbers for all populations are comparable between lines. $n_{TP/Group}$ = 3 - 8 mice.

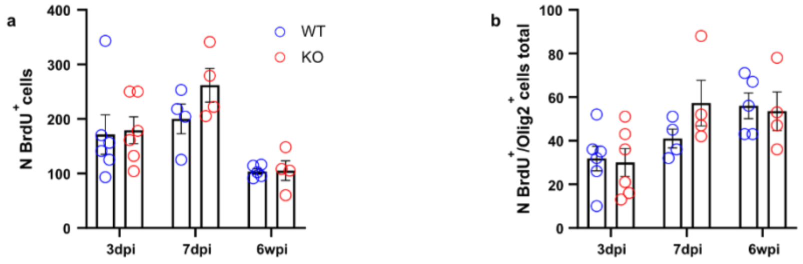


**Figure S6: Quantification of all BrdU⁺ cells and BrdU⁺ OLCs following acute injury in CRHR1$^{\Delta EFGP}$ WT an KO animals. a-b**, Quantification of BrdU⁺ and BrdU⁺/Olig2⁺ cells in 300 µm radius around injury site at 3 dpi, 7 dpi and 6 wpi in *CRHR1$^{\Delta EFGP}$* WT and KO animals showing no significant differences between conditions.

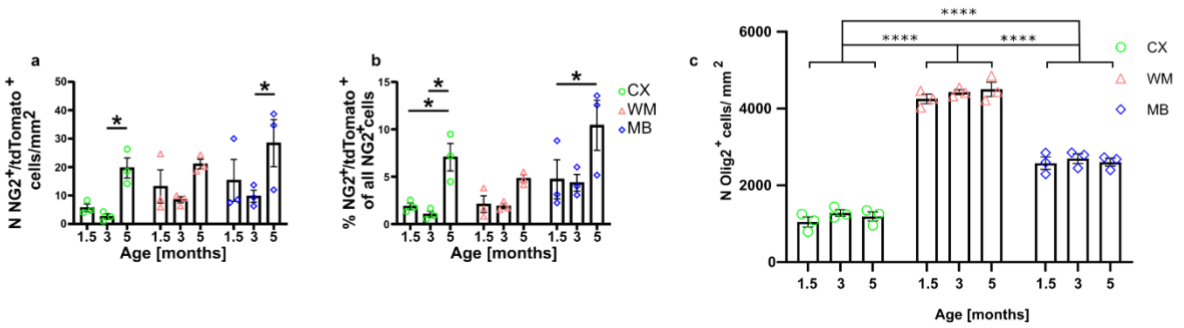


**Figure S7: Quantification of OLC population in non-injured regions a,** Quantification of NG2+/tdTomato+ cells at 1.5, 3 and 5 months of age in CX, WM and MB showing significant increase in CX and MB (Two-way ANOVA: time, $F_{(2, 18)} = 10.34$, $p = 0.001$, $n_{TP} = 3$ mice). **b**, Percentage of NG2+/tdTomato+ of all NG2+ cells showing significant increase over time (Two-way ANOVA: time, $F_{(2, 18)} = 13.46$, $p = 0.003$, $n_{TP} = 3$ mice). **c**, Quantification of Olig2+ cells in CX, WM and MB at 1.5, 3 and 5 months of age shows no increase in the total number of Olig2+ cells. The number of Olig2+ cells are significantly different between regions and highest in WM followed by MB and CX (Two-way ANOVA: region, $F_{(2,18)} = 480.2$, $p < 0.0001$, $n_{TP} = 3$ mice).

**Table S1: Details of used mouse lines**

| Mouse line | Individual Mouse Lines | MGI Allele | MGI Identifier | Description/ Reporting |
|---|---|---|---|---|
| ***CRH-Cre::Ai9*** | *CRH-Cre*<br>*Ai9* | *Crh*$^{tm1(cre)Zjh}$<br>*Gt(ROSA)26Sor*$^{tm9(CAG\text{-}tdTomato)Hze}$ | MGI:4452089<br>MGI:3809523 | tdTomato expression in CRH-expressing cells |
| ***NG2-CreERT2::Ai9*** | *NG2-CreERT2*<br>*Ai9* | *Tg(Cspg4-cre/Esr1*)BAkik*<br>*Gt(ROSA)26Sor*$^{tm9(CAG\text{-}tdTomato)Hze}$ | MGI:4819178<br>MGI:3809523 | tdTomato expression in NG2-expressing cells after induction by tamoxifen |
| ***CRH-Venus*** | *CRH-Venus* | *Crh*$^{tm1.1Ksak}$ | MGI:6144041 | Venus knock-in into *Crh* locus |
| ***CRH-FlpO::NG2-CreERT2::Ai65*** | *CRH-FlpO*<br>*NG2-CreERT2*<br>*Ai65* | *Crh*$^{tm1.1(flpo)Bsab}$<br>*Tg(Cspg4-cre/Esr1*)BAkik*<br>*Gt(ROSA)26Sor*$^{tm65.1(CAG\text{-}tdTomato)Hze}$ | MGI:6116854<br>MGI:4819178<br>MGI:5478743 | tdTomato expression in CRH and NG2 co-expressing cells after induction by tamoxifen |
| ***CRH-FlpO::CRHR1-Cre::Ai65*** | *CRH-FlpO*<br>*CRHR1-Cre*<br>Ai65 | *Crh*$^{tm1.1(flpo)Bsab}$<br>*Crhr1*$^{tm4.1(cre)Jde}$<br>*Gt(ROSA)26Sor*$^{tm65.1(CAG\text{-}tdTomato)Hze}$ | MGI:6116854<br>MGI:6201420<br>MGI:5478743 | tdTomato expression in CRH and CRHR1 co-expressing cells |
| ***CRH***$^{NG2\text{-}cKO}$<br>(*CRH*$^{loxP}$*:: NG2-CreERT2::Ai9*) | *CRH*$^{loxP}$<br>*NG2-CreERT2*<br>*Ai9* | *Crh*$^{tm1.1Jde}$<br>*Tg(Cspg4-cre/Esr1*)BAkik*<br>*Gt(ROSA)26Sor*$^{tm9(CAG\text{-}tdTomato)Hze}$ | MGI:6201415<br>MGI:4819178<br>MGI:3809523 | CRH knockout and tdTomato expression in NG2-expressing cells after induction by tamoxifen; tdTomato allowed quantification of recombination efficiency |
| ***CRHR1***$^{\Delta EGFP}$ | *CRHR1*$^{\Delta EGFP}$ | *Crhr1*$^{tm1Jde}$ | MGI:5294436 | EGFP knock-in into *Crhr1* locus<br>Global/constitutive CRHR1 knockout |
| ***CRHR1-Cre::Tau-LSL-FlpO::Ai9*** | *CRHR1-Cre*<br>*Tau-LSL-FlpO*<br>*Ai9* | *Crhr1*$^{tm4.1(cre)Jde}$<br>*pending*<br>*Gt(ROSA)26Sor*$^{tm9(CAG\text{-}tdTomato)Hze}$ | MGI:6201420<br>pending<br>MGI:3809523 | tdTomato expression in CRHR1-expressing cells; deletion of tdTomato expression in cells with high MAPT expression, enrichment of tdTomato expression in glial CRHR1-expressing cells |
| ***CRHR1-Cre::Sun1-GFP*** | *CRHR1-Cre*<br>*Sun1-GFP* | *Crhr1*$^{tm4.1(cre)Jde}$<br>*Gt(ROSA)26Sor*$^{tm5(CAG\text{-}Sun1/sfGFP)Nat}$ | MGI:6201420<br>MGI:5443817 | GFP expression in nuclear membrane of CRHR1-expressing cells |

| | | | | |
|---|---|---|---|---|
| ***CRH-FlpO::Ai65F::CRHR1-Cre::Sun1-GFP*** | *CRH-FlpO*<br>*Ai65F*<br>*CRHR1-Cre*<br>*Sun1-GFP* | $Crh^{tm1.1(flpo)Bsab}$<br>$Gt(ROSA)26Sor^{tm65.2(CAG\text{-}tdTomato)Hze/J}$<br>$Crhr1^{tm4.1(cre)Jde}$<br>$Gt(ROSA)26Sor^{tm5(CAG\text{-}Sun1/sfGFP)Nat}$ | MGI:6116854<br>MGI:6260212<br>MGI:6201420<br>MGI:5443817 | tdTomato expression in CRH-expressing cells and GFP expression in the nuclear membrane of CRHR1-expressing cells |

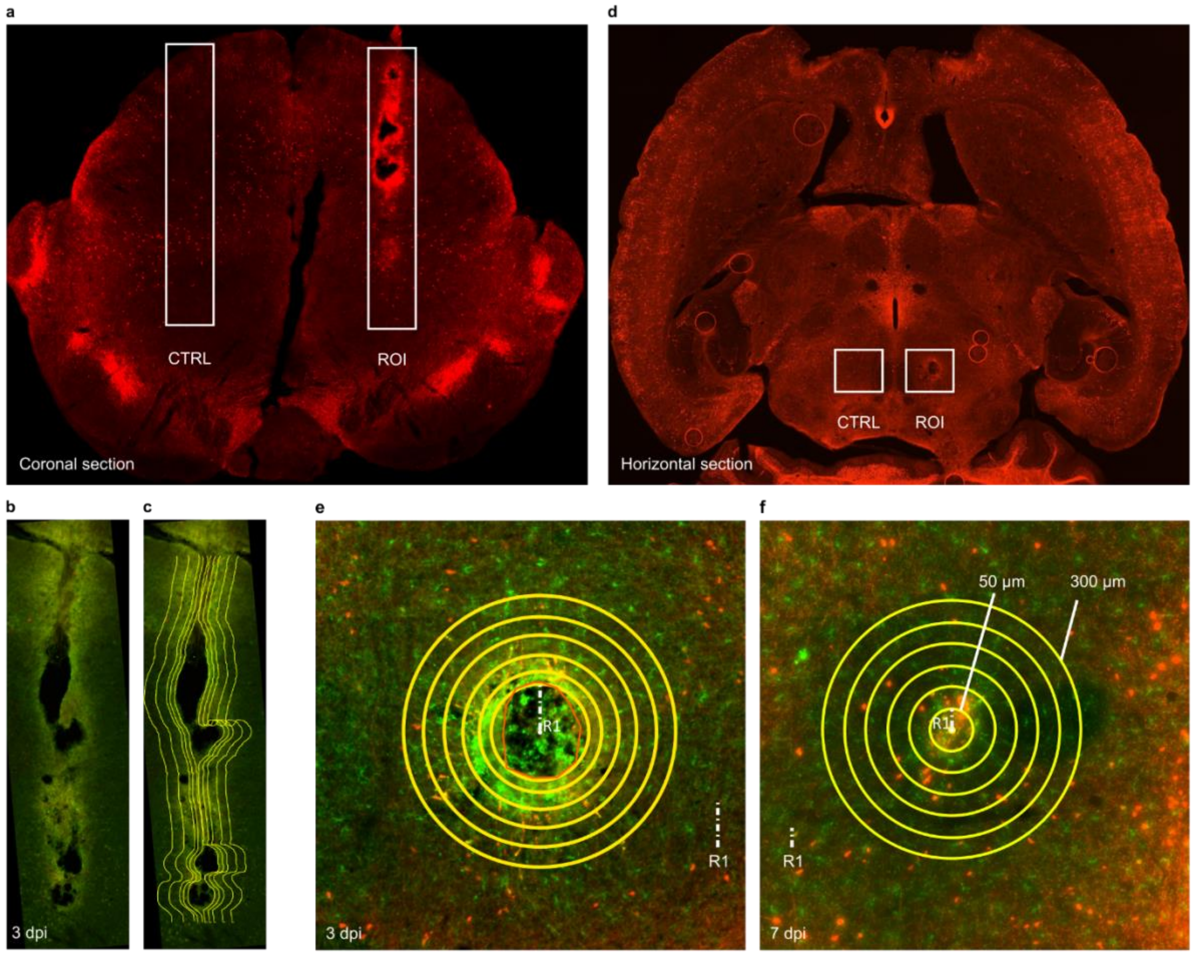


**Method S1: Quantification matrices used for wound analysis. a**, Coronal overview of acute injury in MB of *CRH-Cre::Ai9* mice. White squares: CTRL region and ROI around injury site. **b**, ROI around injury site. **c**, counting matrix generated automatically around center of wound. Subareas from 50 to 300 µm around injury site. **d**, Horizontal overview of acute injury in MB of *CRH-Cre::Ai9* mice. White squares: CTRL region and ROI around injury site. **e, f**, Quantification matrix at 3 dpi (**e**) and 7 dpi (**f**). Matrix consist of 6 circles with radii from 50 to 300 µm.
